# Optimism and pessimism in optimised replay

**DOI:** 10.1101/2021.04.27.441454

**Authors:** Georgy Antonov, Christopher Gagne, Eran Eldar, Peter Dayan

## Abstract

The replay of task-relevant trajectories is known to contribute to memory consolidation and improved task performance. A wide variety of experimental data show that the content of replayed sequences is highly specific and can be modulated by reward as well as other prominent task variables. However, the rules governing the choice of sequences to be replayed still remain poorly understood. One recent theoretical suggestion is that the prioritization of replay experiences in decision-making problems is based on their effect on the choice of action. We show that this implies that subjects should replay sub-optimal actions that they dysfunctionally choose rather than optimal ones, when, by being forgetful, they experience large amounts of uncertainty in their internal models of the world. We use this to account for recent experimental data demonstrating exactly pessimal replay, fitting model parameters to the individual subjects’ choices.

## Introduction

During periods of quiet restfulness and sleep, when humans and other animals are not actively engaged in calculating or executing the immediate solutions to tasks, the brain is nevertheless not quiet. Rather it entertains a seething foment of activity. The nature of this activity has been most clearly elucidated in the hippocampus of rodents, since decoding the spatial codes reported by large populations of simultaneously recorded place cells [1, 2] reveals ordered patterns. Rodents apparently re-imagine places and trajectories that they recently visited (‘replay’) [3, 4, 5, 6], or might visit in the future (‘preplay’) [6, 7, 8, 9, 10, 11, 12, 13], or are associated with unusually large amounts of reward [14, 15, 16]. However, replay is not only associated with the hippocampus; there is also a complex semantic and temporal coupling with dynamical states in the cortex [17, 18, 19, 20, 21, 22, 23].

In humans, the patterns of neural engagement during these restful periods have historically been classified in such terms as default mode or task-negative activity [24]. This activity has been of great value in elucidating functional connectivity in the brain [25, 26, 27]; however, its information content had for a long time been somewhat obscure. Recently, though, decoding neural signals from magnetoencephalographic (MEG) recordings in specific time periods associated with the solution of carefully designed cognitive tasks, has revealed contentful replay and preplay (for convenience, we will generally refer to both simply as ‘replay’) that bears some resemblance to the rodent recordings [28, 29, 30, 31, 32].

The obvious question is what computational roles, if any, are played by these informationally-rich signals. It is known that disrupting replay in rodents leads to deficits in a variety of tasks [33, 34, 35, 19, 36], and there are various theoretical ideas about its associated functions. Although the notions are not completely accepted, it has been suggested that the brain uses off-line activity to build forms of inverse models – index extension and maintenance in the context of memory consolidation [37]; recognition models in the case of unsupervised learning [38], and off-line planning in the context of decision-making [39, 40].

While appealing, these various suggestions concern replay in general, and have not explained the micro-structure of which pattern is replayed when. One particularly promising idea for this in the area of decision-making is that granular choices of replay experiences are optimized for off-line planning [41]. The notion, which marries two venerable suggestions in reinforcement learning (RL) [42]: DYNA [39] and prioritized sweeping [43], is that each replayed experience changes the model-free value of an action in order to maximize the utility of the animal’s ensuing behaviour. It was shown that the resulting optimal choice of experience balances two forces: need, which quantifies the expected frequency with which the state involved in the experience is encountered, and gain, which quantifies the benefit of the change to the behaviour at that state occasioned by replaying the experience. This idea explained a wealth of replay phenomena in rodents.

Applying these optimizing ideas to humans has been hard, since, until recently the micro-structure of replay in humans had not been assessed. Liu *et al*. 2020 [32] offered one compelling test which well followed Mattar and Daw 2018 [41]. By contrast, in a simple planning task, Eldar *et al*. 2020 [31] showed an unexpected form of efficacious replay in humans that, on the surface, seemed only partially to align with this theory. In this task, subjects varied in the extent to which their decisions reflected the utilisation of a model of how the task was structured. The more model-based (MB) they were, the more they engaged in replay during inter-trial interval periods, in a way that appeared helpful for their behaviour. Strikingly, though, the replay was apparently pessimized – that is, subjects preferred to replay *bad* choices, which were then deprecated in the future.

In this paper, we consider this characteristic of replay, examining it from the perspective of optimality. We show that favouring bad choices is in fact appropriate in the face of substantial uncertainty about the transition structure of the environment – a form of uncertainty that arises, for instance, from forgetting. Moreover, we consider the costs and benefits of replay on task performance in the light of subjects’ subjective, and potentially deviant, knowledge of the task. Although, to be concrete, we focus on the task studied by Eldar *et al*. 2020 [31], the issues we consider are of general importance.

## Results

### Preamble

In the study of Eldar *et al*. 2020 [31], human subjects acted in a carefully designed planning task (Fig 1A). Each state was associated with an image that was normally seen by the subjects. The subjects started each trial in a pseudo-random state and were required to choose a move among the 4 possible directions: up, down, left or right (based on the toroidal connectivity shown). In most cases, they were then shown an image associated with the new state according to the chosen move and received a reward associated with that image (this was not displayed; however, the subjects had been extensively taught about the associations between images and reward). Some trials only allowed single moves and others required subjects to make an additional second move which provided a second reward from the final state. In most of these 2-move trials, in order to obtain maximal total reward, the subjects had to select a first move that would have been sub-optimal had it been a 1-move trial.

**Fig. 1.**
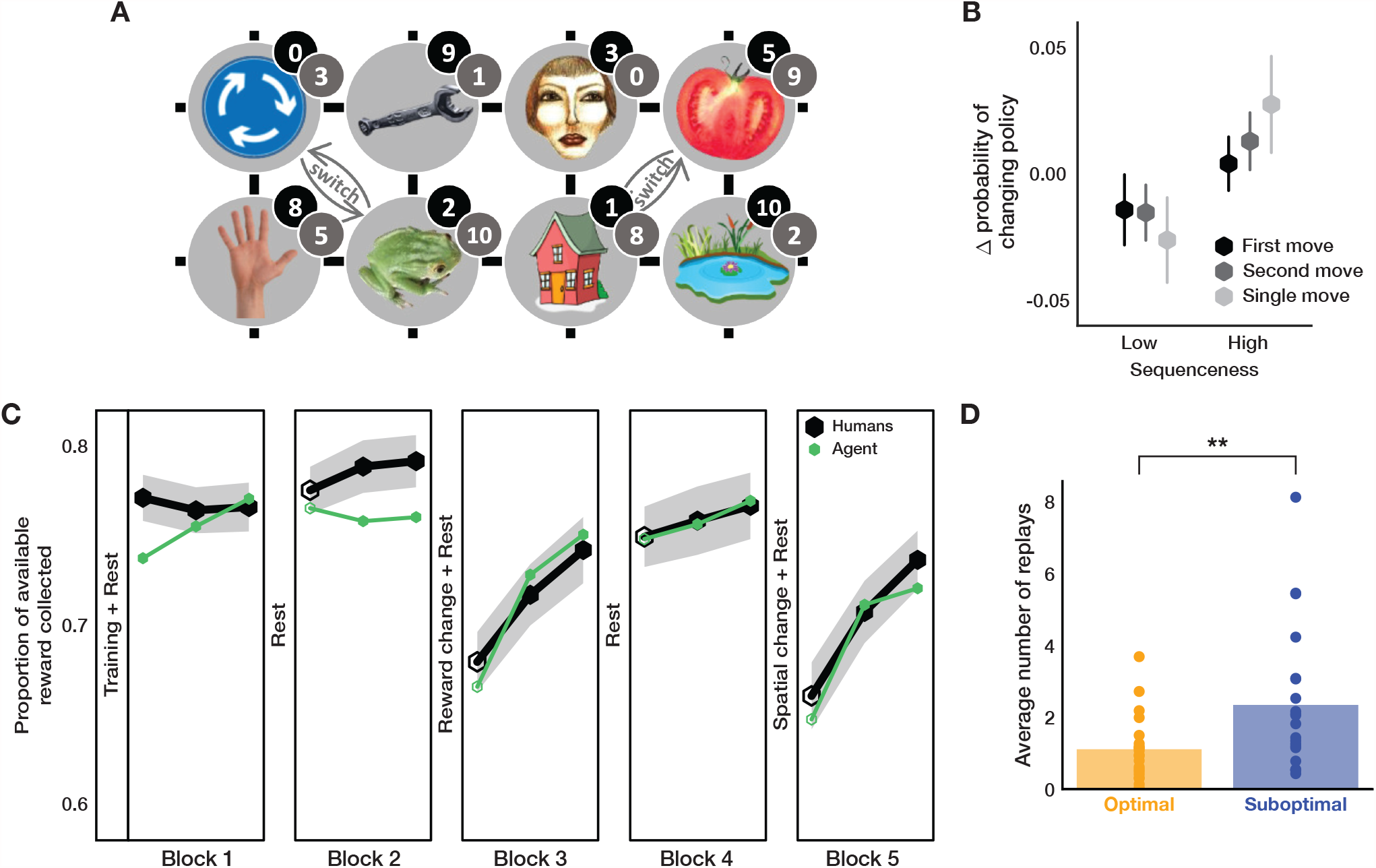
Task Structure and Replay Modelling. (A) Structure of the state-space. Numbers in black and grey circles denote the number of reward points associated with that state respectively pre- and post-the reward association change between blocks 2 and 3. Grey arrows show the spatial re-arrangement that took place between blocks 4 and 5. (B) Change in the probability of choosing a different move when in the same state as a function of sequenceness of the just-experienced transitions measured from the MEG data in subjects with non-negligible sequenceness (*n* = 25). High sequenceness was defined as above median and low sequenceness as below median. Analysis of correlation between the decoded sequenceness and probability of policy change indicated a significant dependency (Spearman correlation, *M* = 0.04, *SEM* = 0.02, *p* = 0.04, Bootstrap test). Vertical lines show SEM. (C) Performance of the human subjects and the agent with parameters fit to the individual subjects. Unfilled hexagons show epochs which contained trials without feedback. Shaded area shows SEM. (D) Pessimism bias in the replay choices of human subjects for which our model predicted sufficient replay (*n* = 21) as reflected in the average number of replays of recent sub-optimal and optimal transitions at the end of each trial (sub-optimal vs optimal, Wilcoxon rank-sum test, *W* = 2.60, *p* = 0.009). ** *p <* 0.01.

The subjects were not aware of the spatial arrangement of the state space, and thus had to learn about it by trial and error (which they did in the training phase that preceded the main task, see Methods for details). In order to collect additional data on subjects’ knowledge of the state space, no feedback was provided about which state was reached after performing an action in the first 12 trials of blocks two through five. This feedback was provided in the remaining 42 trials to allow ongoing learning in the face, for instance, of forgetting. After two blocks of trials, the subjects were taught a new set of associations between images and rewards; similarly, before the final block they were informed about a pair of re-arrangements in the transition structure of the state space (involving swapping the locations of two pairs of images).

In order to achieve high performance, the subjects had to a) adjust their choices according to whether the trial allowed 1 or 2 moves; and b) adapt to the introduced changes in the environment. Eldar *et al*. 2020 [31] calculated an individual flexibility (IF) index from the former adjustment, and showed that this measure correlated significantly with how well the subjects adapted to the changes in the environment (and with their ability to draw the two different state spaces after performing all the trials, as well as to perform 2-move trials in the absence of feedback about the result of the first move). The more flexible subjects therefore presumably utilised a model of the environment to plan and re-evaluate their choices accurately.

Eldar *et al*. 2020 [31] investigated replay by decoding MEG data to reveal which images (i.e., states) subjects were contemplating during various task epochs. They exploited the so-called sequenceness analysis [28] to show that, in subjects with high IF, the order of contemplation of states in the inter-trial intervals following the outcome of a move revealed the replay of recently visited transitions (as opposed to the less flexible, presumably model-free (MF), subjects); it was notable that the transitions they preferred to replay (as measured by high sequenceness) mostly led to sub-optimal outcomes. Nevertheless, those subjects clearly benefited from what we call pessimized replay, for after the replay of sub-optimal actions they were, correctly, less likely to choose these (Fig 1B) when faced with the same selection of choices later on in the task.

In Eldar *et al*. 2020 [31], and largely following conventional suggestions [44, 45, 46, 47], subjects’ choices were modelled with a hybrid MF/MB algorithm. This fit the data better than algorithms that relied on either pure MF or MB learning strategies. However, the proposed model did not account for replay and therefore could not explain the preference for particular replays or their effect on choice.

Therefore, to gain further insights into the mechanisms that underpinned replay choices of human subjects (as well as their effect on behaviour), we constructed an agent that made purely MF choices, but whose MF values were adjusted by a form of MB replay [41] that was optimal according to the agent’s forgetful model of the task. The agent was therefore able to adapt its decision strategy flexibly by controlling the amount of influence maintained by (subjectively optimal) MB information over MF values. We simulated this agent in the same behavioural task with the free parameters fit to the subjects (Fig 1C), and examined the resulting replay preferences.

### Modelling of Subjects’ Choices in the Behavioural Task

To model replay in the behavioural task, we used a DYNA-like agent [39] which learns on-line by observing the consequences of its actions, as well as off-line in the inter-trial intervals by means of generative replay (Fig 2A). On-line learning is used to update a set of MF *Q*-values [48] which determine the agent’s choices through a softmax policy, as well as to (re)learn a model of the environment (i.e., transition probabilities). During off-line periods, the agent uses its model of the transition structure of the environment to estimate *Q*-values (denoted as 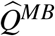), and then evaluates these MB estimates for the potential improvements to MF policy that the agent uses to make decisions. This process of evaluation and improvement is the key difference between our model and that of Eldar *et al*. 2020 [31]: instead of having MB quantities affecting the agent’s choices directly, they only did so by informing optimized replay [41] that provided additional training for the MF values.

**Fig. 2.**
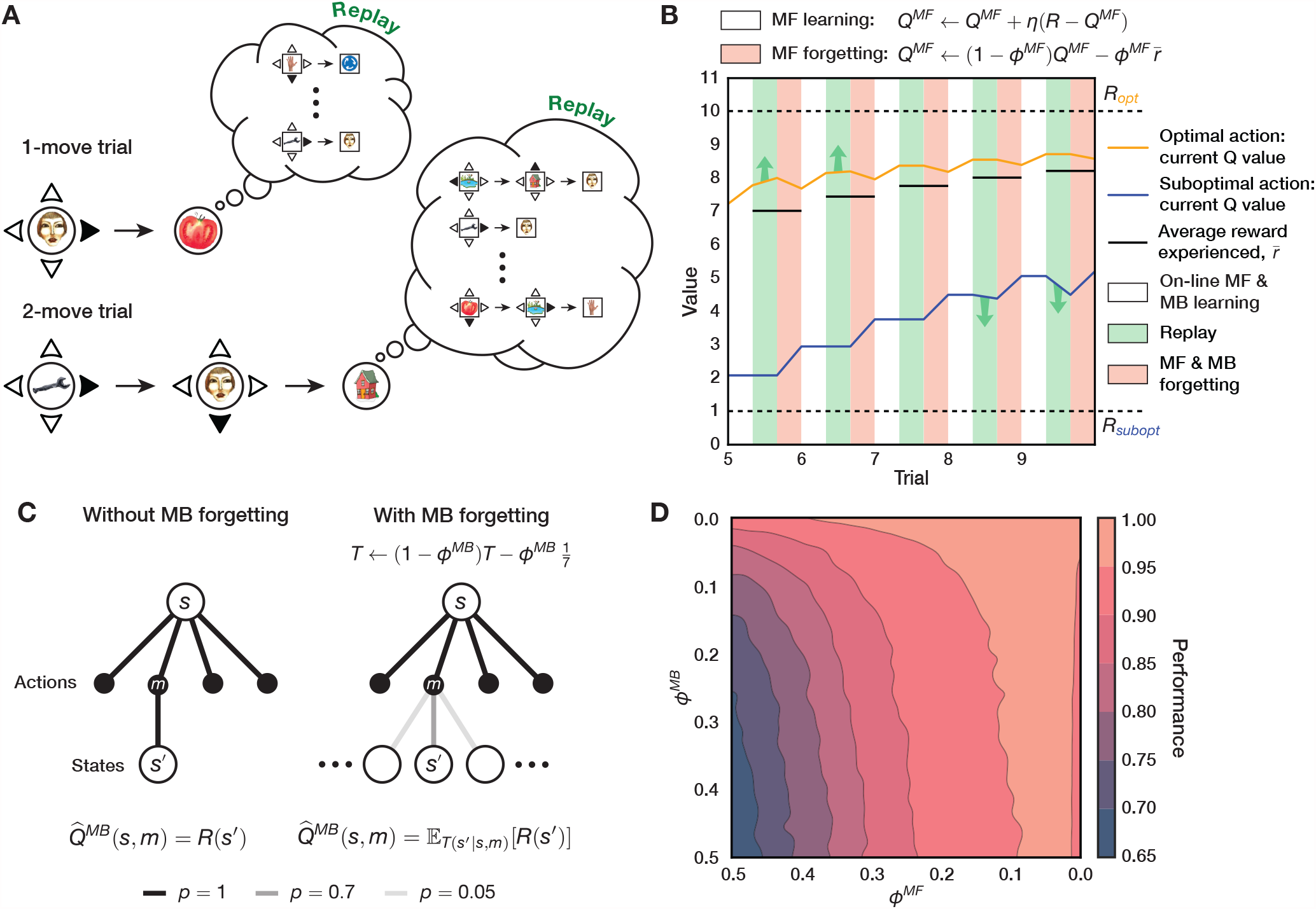
Algorithm Description and the Effects of Replay and Forgetting on Model Performance. (A) Schematic illustration of the algorithm in the behavioural task. Upon completing each trial, the algorithm uses its knowledge of the transition structure of the environment to replay the possible outcomes. Note that in 1-move trials the algorithm replays only single moves, while in 2-move trials it considers both single and coupled moves (thus optimizing this choice). (B) Effect of MF forgetting and replay on MF *Q*-values. After acting and learning on-line towards true reward *R* (white blocks; controlled by learning rate, *η*), the algorithm learns off-line by means of replay (green blocks). Immediately after each replay bout, the algorithm forgets its MF *Q*-values towards the average reward experienced from the beginning of the task (red blocks; controlled by MF forgetting rate, *ϕ* ^*MF*^). Note that after trials 5 and 6, the agent chooses to replay the objectively optimal action, whereas after trials 8 and 9 it replays the objectively sub-optimal action. (C) Left: without MB forgetting, the algorithm’s estimate of reward obtained for a given move corresponds to the true reward function. Right: with MB forgetting (controlled by MB forgetting rate, *ϕ* ^*MB*^), the algorithm’s estimate of reward becomes an expectation of the reward function under its state transition model. The state-transition model’s probabilities for the transitions are shown as translucent lines. (D) Steady-state performance (proportion of available reward obtained) of the algorithm in the behavioural task as a function of MF forgetting, *ϕ* ^*MF*^, and MB forgetting, *ϕ* ^*MB*^. Note how the agent still achieves high performance with substantial MF forgetting (high *ϕ* ^*MF*^) when its state-transition model accurately represents the transition probabilities (low *ϕ* ^*MB*^).

Unlike a typical DYNA agent, or indeed the suggestion from Mattar and Daw 2018 [41], our algorithm performs a full model evaluation, and therefore MF *Q*-values are updated in a supervised manner towards the model-generated 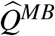 values. It is crucial to note that subjects were found by Eldar *et al*. 2020 [31] (and also in our own model fitting) to be notably forgetful. Therefore, although the task is deterministic, replay can help (and hurt) since the seemingly omniscient MB updates may in fact be useless, or even worse, harmful. To avoid extensive training of MF values, in the light of the potential harm an inaccurate MB system can accomplish, the agent thus only engages in replay as long as the potential MB updates are estimated to be sufficiently gainful (see below); this is controlled by a replay gain threshold, which was a free parameter that we fit to each individual subject.

Similarly to the best-fitting model in Eldar *et al*. 2020 [31], after experiencing a move, our agent forgets both MF *Q*-values and the state-transition model (for all allowable transitions). However, unlike that account, over the course of forgetting, MF *Q*-values tend towards the average reward the agent has experienced since the beginning of the task (Fig 2B), as opposed to tending towards what was a fixed subject-specific parameter. Insofar as the agent improves over the course of the task, the average reward it obtains increases with each trial. MF *Q*-values for sub-optimal actions, therefore, tend to rise towards this average experienced reward; MF *Q*-values for optimal actions, on the other hand, become devalued, as the agent is prone occasionally to choose sub-optimal actions due to its non-deterministic policy. In other words, because of MF forgetting, the agent forgets how good the optimal actions are and how bad the sub-optimal actions are. Similarly, because of MB forgetting, the agent gradually forgets what is specific about particular transitions, progressively assuming a uniform distribution over the potential states to which it can transition (Fig 2C). The agent therefore becomes uncertain over time about the consequences of actions it rarely experiences.

The two forgetting mechanisms significantly influence the agent’s behaviour – MF forgetting effectively decreases the value of each state by infusing the agent’s policy with randomness, whereas MB forgetting confuses the agent with respect to the individual action outcomes. From an optimality perspective, the question is then what and if the agent should replay at all, given the imperfect knowledge of the world and a forgetful MF policy. We find that at high MF forgetting, replay confers a noticeable performance advantage to the agent provided that MB forgetting is mild (as can be seen from the curvature of the contour lines in Fig 2D). This means that despite moderate uncertainty in the transition structure, the agent is still able to improve its MF policy and increase the obtained reward rate.

We then analysed the replay choices of human participants which, according to our model prediction (with the free parameters of our agent fit to data from individual subjects), engaged in replay. This revealed a significant preference to replay actions that led to sub-optimal outcomes (Fig 1D). We therefore considered the parameter regimes in our model that led the agent to make such pessimal choices, and whether the subjects’ apparent preference to replay sub-optimal actions was formally beneficial for improving their policies.

### Exploration of Parameter Regimes

The analysis in Mattar and Daw 2018 [41] suggests that two critical factors, need and gain, should jointly determine the ordering of replay by which an (in their case, accurate) MB system should train an MF controller. Need quantifies the extent to which a state is expected to be visited according to the agent’s policy and transition dynamics of the environment. It is closely related to the successor representation across possible start states, which is itself a prediction of discounted future state occupancies [49]. Heterogeneity in need would come from biases in the initial states on each trial (5 of the 8 were more common; but which 5 changed after blocks 2 and 4) and the contribution of subject’s preferences for the first move on 2-move trials. However, we expected the heterogeneity itself to be modest and potentially hard for subjects to track, and so made the approximation that need was the same for all states.

Gain quantifies the expected local benefit at a state from the change to the policy that would be engendered by a replay. Importantly, gain only accrues when the behavioural policy changes. Thus, one reason that the replay of sub-optimal actions is favoured is that replays that strengthen an already apparently optimal action will not be considered very gainful – so to the extent that the agent already chooses the best action, it will have little reason to replay it. A second reason comes from considering why continuing learning is necessary in this context anyhow – i.e., forgetting of the MF *Q*-values. Since the agent learns on-line as well as off-line, it will have more opportunities to learn about optimal actions without replay. Conversely, the values of sub-optimal actions are forgotten without this compensation, and so can potentially benefit from off-line replay.

We follow Mattar and Daw 2018 [41] in assuming that the agent computes gain optimally. However, this optimal computation is conducted on the basis of the agent’s subjective model of the task, which will be imperfect given forgetfulness. Thus, a choice to replay a particular transition might seem to be suitably gainful, and so selected by the agent, but would actually be deleterious, damaging the agent’s ability to collect reward. In this section, we explore this tension.

In Fig 3A, we show the estimated gain for an objectively optimal and sub-optimal action in an example simulation with only two actions available to the agent (see Methods for details). The bar plot in Fig 3B illustrates the ‘centering’ effect MF forgetting has on sub-optimal MF *Q*-values; this effect, however, is not symmetric. As the agent learns the optimal policy, the average reward it experiences becomes increasingly similar to the average reward obtainable from optimal actions in the environment. As a result, there is little room for MF *Q*-values for optimal actions to change through forgetting; on the other hand, MF *Q*-values for sub-optimal actions are forgotten towards what is near to the value of optimal actions to a much more substantial degree. Thus forgetting is optimistic in a way that favours replay of sub-optimal actions.

**Fig. 3.**
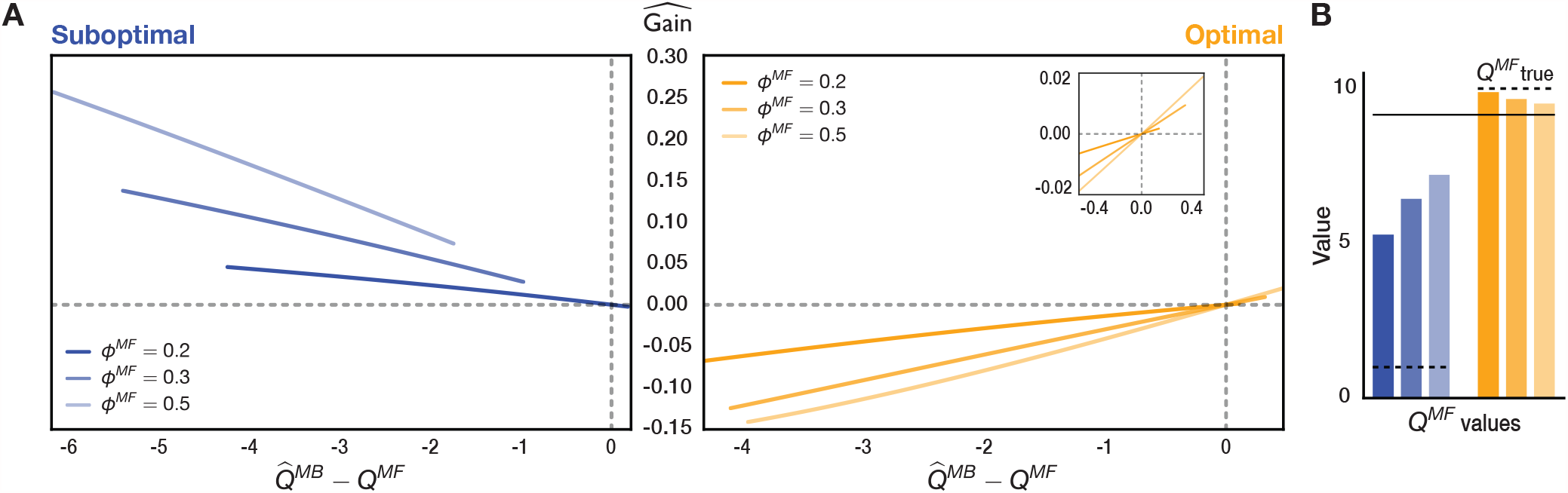
How MF Forgetting Influences Gain Estimation. (A) Estimated gain as a function of the difference between the agent’s current MF *Q*-value and the model-estimated MB *Q*-value, 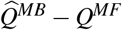, for varying degrees of MF forgetting, *ϕ* ^*MF*^. The dashed grey lines show the x- and y-intercepts. Note that the estimated gain is negative whenever the model-generated 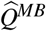 estimates are worse than the current MF *Q*-values. (B) Current MF *Q*-values for the optimal and sub-optimal actions with varying MF forgetting rate, coloured in the same way as above. The horizontal solid black bar is the average reward experienced so far, towards which MF values tend. The true *Q*-value for each action is shown in dashed black.

Objectively sub-optimal actions that our agent replays therefore mostly have negative temporal difference (TD) errors. Such pessimized replay reminds the agent that sub-optimal actions, according to its model, are actually worse than predicted by the current MF policy.

There is an additional, subtle, aspect of forgetting that decreases both the objective and subjective benefit of replay of what are objectively optimal actions (provided the agent correctly estimates such actions to be optimal). For the agent to benefit from this replay, its state-transition model must be appropriately accurate as to generate MB *Q*-values for optimal actions that are better than their current MF *Q*-values. Forgetting in the model of the effects of actions (i.e., transition probabilities) will tend to homogenize the expected values of the actions – and this exerts a particular toll on the actions that are in fact the best. Conversely, forgetting of the effects of sub-optimal actions if anything makes them more prone to be replayed. This is because any sub-optimal action considered for replay will have positive estimated gain as long as MB *Q*-value for that action (as estimated by the agent’s state-transition model) is less than its current MF *Q*-value. Due to MF forgetting, the agent’s MF *Q*-values for sub-optimal actions rise above the actual average reward of the environment, and therefore, even if the state-transition model is uniform – which is the limit of complete forgetting of the transition matrix, the agent is still able to generate MB *Q*-values for sub-optimal actions that are less than their current MF *Q*-values, and use those in replay to improve the MF policy.

To examine these, we quantified uncertainty in the agent’s state-transition model, for every state *s* and action *m* considered for replay, using the standard Shannon entropy [50]:

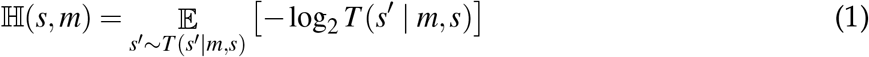

over the potential states to which the agent could transition. Similarly, the agent’s uncertainty about 2-move transitions considered for replay was computed as the joint entropy of the two transitions (equation 23). For any state *s* and action *m* we will henceforth refer to ℍ (*s, m*) as the action entropy (importantly, it is not the overall model entropy since the agent can be more or less uncertain about particular transitions). If an action that is considered for replay has high action entropy, the estimated MB *Q*-value of that action is corrupted by the possibility of transitioning into multiple states (Fig 4A, left); in fact, maximal action entropy corresponds to a uniform policy. For an action with low action entropy, on the other hand, the agent is able to estimate the MB *Q*-value of that action more faithfully (Fig 4A, right).

**Fig. 4.**
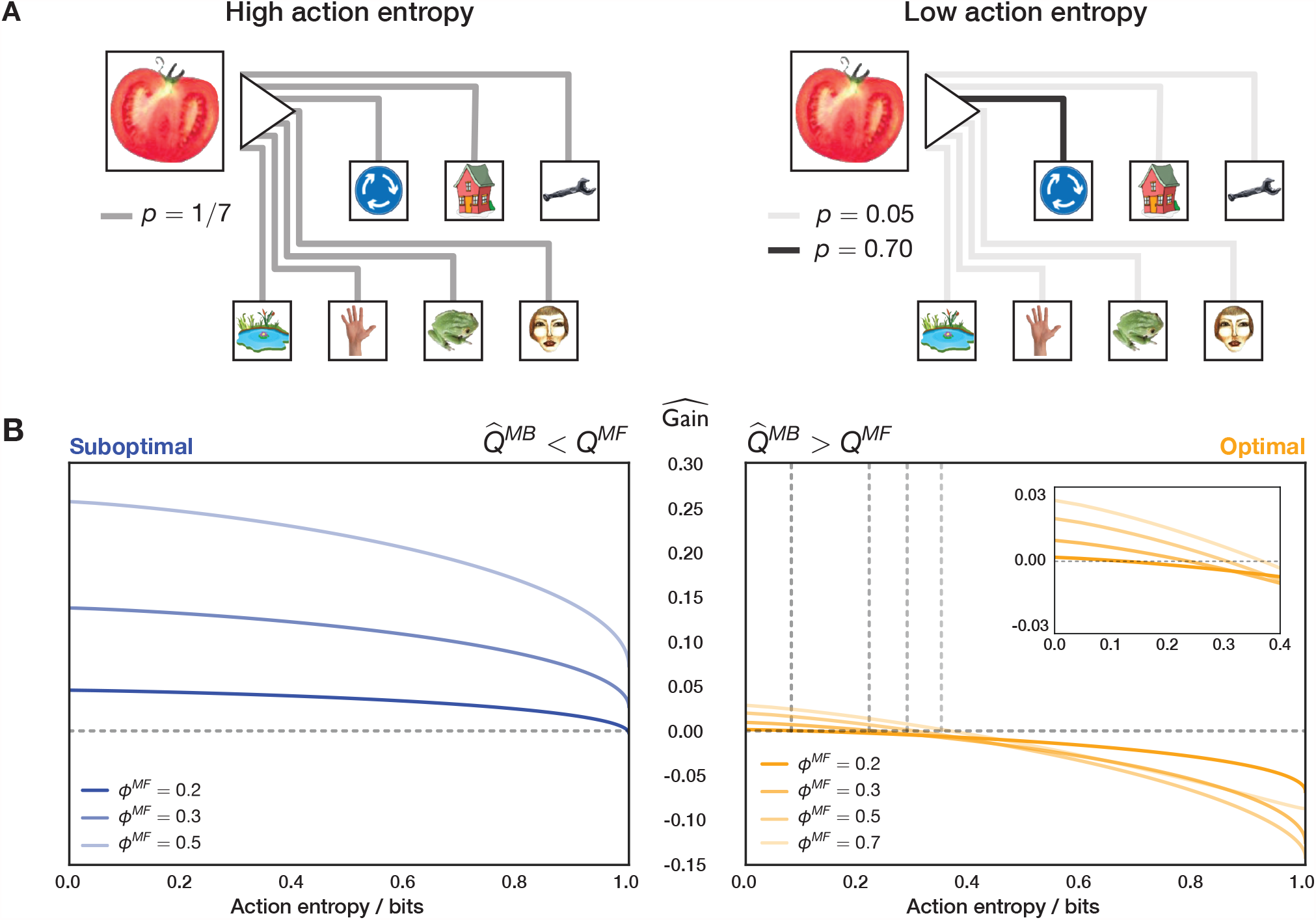
Action Entropy Limits Estimated Gain. (A) Left: under high action entropy, the distribution over the potential states to which the agent can transition given the current state and a chosen action is close to uniform. Right: under low action entropy, the agent is more certain about the state to which a chosen action will transition it. (B) Left: for an objectively sub-optimal action, the gain is positive throughout most action entropy values. Right: for an objectively optimal action, the gain becomes positive only when the state-transition model is sufficiently accurate. With heavier MF forgetting (higher *ϕ* ^*MF*^), however, the intercept shifts such that the agent is able to benefit from a less accurate model (grey dashed lines show the x- and y-intercepts). The inset magnifies the estimated gain for the optimal action. Moreover, note how the magnitude of the estimated gain for an objectively optimal action is lower than that of a sub-optimal one, which is additionally influenced by the asymmetry of MF forgetting and on-line learning.

Thus, we examined how the estimated gain of objectively optimal and sub-optimal actions is determined by action entropy (Fig 4B). Indeed, we found that the estimated gain for optimal actions was positive only at low action entropy values (Fig 4B, right; see also S1 Fig), hence confirming that uncertainty in the agent’s state-transition model significantly limited its ability to benefit from the replay of optimal actions. By contrast, the estimated gain for sub-optimal actions was positive for a wider range of action entropy values, compared to that for optimal actions (Fig 4B, left; see also S1 Fig).

Of course, optimal actions are performed more frequently, thus occasioning more learning. This implies that the state-transition model will be more accurate for the effects of those actions than for sub-optimal actions, which will partly counteract the effect evident in Fig 4. On the other hand, as discussed before, the estimated gain for optimal actions will be, in general, lower, precisely because of on-line learning as a result of the agent frequenting such actions.

An additional relevant observation is that action entropy (due to MB forgetting), together with MF forgetting, determines the balance between MF and MB strategies to which the agent apparently resorts. This can be seen from the increasing x-intercept as a function of the strength of MF forgetting for optimal actions (vertical dashed lines in Fig 4B, right). In the case of weak MF forgetting, the agent can only benefit from the replay of optimal actions inasmuch as the state-transition model is sufficiently accurate – which requires action entropy values to be low.

As MF forgetting becomes more severe, the agent can think itself to benefit from replay extracted from a worse state-transition model. Thus, we identify a range of parameter regimes which can lead agents to find replay subjectively beneficial and, therefore, allocate more influence to the MB system.

### Fitting Actual Subjects

We fit the free parameters of our model to the individual subject choices from the study of Eldar *et al*. 2020 [31], striving to keep as close as possible to the experimental conditions, for instance by treating the algorithm’s adaptations to the image-reward association changes and the spatial re-arrangement in the same way as in the original study (see Methods for details). First, we examined whether our model correctly captured the varying degree of decision flexibility that was observed across subjects. Indeed, we found that the simulated IF values, as predicted by our agent with subject-tailored parameters, correlated significantly with the behavioural IF values (S3 Fig A). Moreover, our agent predicted that some subjects would be engaging in potentially measurable replay (an average of more than 0.5 replays per trial, *n* = 21), and hence use MB knowledge when instructing their MF policies to make decisions. Indeed, we found that our agent predicted those subjects to be significantly less ignorant about the transition probabilities at the end of the training trials (S8 Fig), thus indicating the accumulation of MB knowledge. Further, those same subjects had significantly higher simulated IF values relative to the subjects for whom the agent did not predict sufficient replay – which is in line with the observation of Eldar *et al*. 2020 [31] that subjects with higher IF had higher MEG sequenceness following ‘surprising’ (measured by individually-fit state prediction errors) trial outcomes.

We then examined the replay patterns of those subjects, which we refer to as model-informed (MI) subjects, when modelled by our agent. There is an important technical difficulty in doing this exactly: our agent was modelled in an on-policy manner – i.e., making choices and performing replays based on its subjective gain, which, because of stochasticity, might not emulate those of the subjects, even if our model exactly captured the mechanisms governing choice and replay in the subjects. However, we can still hope for general illumination from the agent’s behaviour.

In Fig 5A, we show an example move in a 1-move trial which was predicted by our agent with parameters fit to an MI subject. This example is useful for demonstrating how the agent’s forgetful state of knowledge (as discussed in the previous section) led it to prioritise the replay of certain experiences, and whether those choices were objectively optimal.

**Fig. 5.**
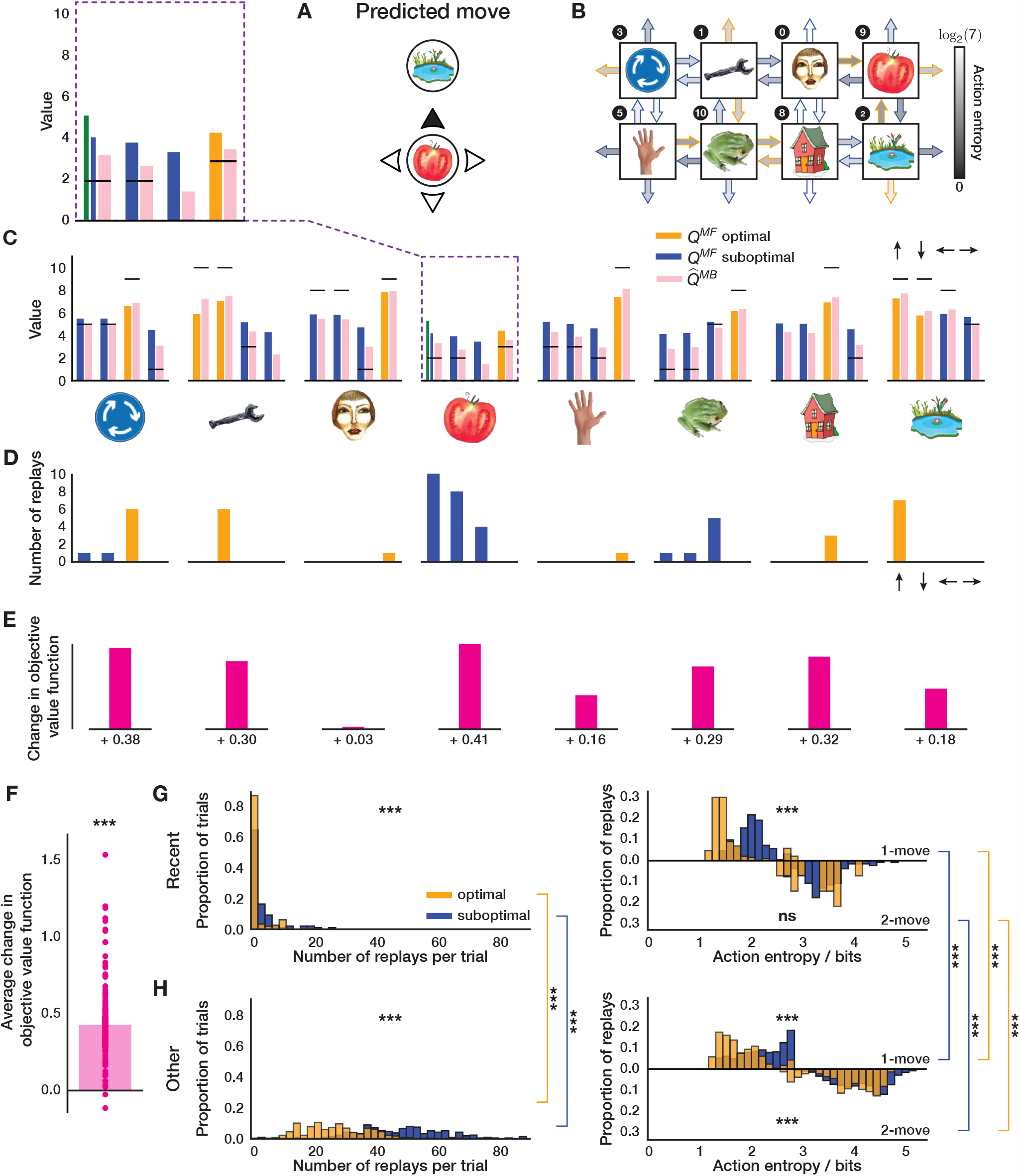
Epistemology of Replay. (A) An example move predicted by our agent with subject-specific parameters. (B) State-transition model of the agent after executing the move in (A) and the associated action entropy values. Objectively optimal actions are shown as arrows with orange outlines; sub-optimal – with blue outlines. Continued on next page. (C) State of MF and MB knowledge of the agent. The arrows above the rightmost bar plot indicate the directions of the corresponding actions in each plot. The horizontal black lines represent the true reward obtainable for each action. The agent’s knowledge at the state where the trial began is highlighted in a purple dashed box and is additionally magnified above. The green bar for the MF *Q*-value that corresponds to the predicted move in (A) shows what the agent knew before executing the move, and the neighbouring blue bar – what the agent has learnt online after executing the move (note that the agent always learnt online towards the true reward). (D) Replay choices of the agent. (E) Changes in the objective value function (relative to the true obtainable reward) of each state as a result of the replay in (D), not drawn to scale. (F) Same as in (D) but across the entire experiment and averaged over all states. (G-H) Average replay statistics over the entire experiment. (G) Just-experienced transitions; (H) Other transitions. First column: proportion of sub-optimal and optimal trials in which objectively sub-optimal or optimal action(s) were replayed. Second column: proportion of action entropy values at which the replays were executed. Upper and lower y-axes show the action entropy distribution for 1-move and 2-move trials respectively. *** *p <* 0.001, ns: not significant.

In this particular example, the agent chose a sub-optimal move. The agent’s state of knowledge of 1-move transitions before and after executing the same move is shown in Fig 5B and c. Note how the agent is less ignorant, on average, about the outcomes of 1-move optimal actions, as opposed to sub-optimal ones (Fig 5B). The agent chose a sub-optimal move because the MF *Q*-value for that move had been forgotten to an extent that the agent’s subjective knowledge incorrectly indicated it to be optimal (Fig 5C). After learning online that the chosen move was worse than predicted (Fig 5C, compare green and blue bars), the agent replayed that action at the end of the trial to incorporate the estimated MB *Q*-value for that action, as generated by the agent’s state-transition model (Fig 5C, pink bar), into its MF policy (Fig 5D). In this case, the MB knowledge that the agent decided to incorporate into its MF policy was more accurate as regards the true *Q*-values; such replay, therefore, made the agent less likely to re-choose the same sub-optimal move in the following trials. Note that even though the actual move (‘up’) was replayed ten times, the action (‘down’) that was replayed eight times led to the same state, and would be indistinguishable in the MEG signal as measured by sequenceness. Therefore, in this example, the agent preferentially engaged in the replay of the recent sub-optimal single move experience.

As just discussed, in addition to replaying the just-experienced transition, the agent also engaged in the replay of other transitions. Indeed, the agent was not restricted in which transitions to replay – it chose to replay actions based solely on whether the magnitude of the estimated gain of each possible action is greater than a subject-specific gain threshold (see Methods for details). We were, therefore, able to see a much richer picture of the replay of all allowable transitions, in addition to the just-experienced ones. In the given example, it is easy to see how the agent estimated its replay choices to be gainful: because of the relatively strong MF forgetting and a sufficiently accurate state-transition model. To demonstrate a slightly different parameter regime, we also additionally show an example move predicted by our agent for another MI subject (S4 Fig). In that case, the agent’s MF and MB policies were more accurate; however, our parameter estimates indicated that the subject’s gain threshold for initiating replay was set lower, and hence the agent still engaged in replay, even though the gain it estimated was apparently minute.

To quantify the objective benefit of replay for the subject shown in Fig 5, we examined how the agent’s objective value function (relative to the true obtainable reward) at each state changed due to the replay at the end of this trial, which is a direct measure of the change in the expected reward the agent can obtain. We found that for each state where the replay occurred, the objective value function of that state increased (Fig 5E). To see whether such value function improvements held across the entire session, we examined the average trial-wise statistics for this subject (Fig 5F). We found that, on average, the replay after each trial (both 1-move and 2-move) improved the agent’s objective value function by 0.43 reward points (1-sample 2-tailed t-test, *t* = 31.3, *p* ≪ 0.0001).

We next looked at the average trial-wise replay statistics of the subject as predicted by our agent (Fig 5G, H). If we consider solely the just-experienced transitions (Fig 5G), we found that there were significantly more replays of sub-optimal actions per trial (Wilcoxon rank-sum test, *W* = 4.05, *p* = 5.15 · 10^−5^). Moreover, the replay of sub-optimal single actions was at significantly higher action entropy values (Wilcoxon rank-sum test, *W* = 18.3, *p* ≪ 0.0001), which is what one would expect given that the transitions that led to optimal actions were experienced more frequently. Since some optimal 1-move actions corresponded to sub-optimal first moves in 2-move trials, the agent received an additional on-line training about the latter transitions, and we therefore found no significant difference in action entropy values at which coupled optimal and sub-optimal actions were replayed (Wilcoxon rank-sum test, *W* = −0.07, *p* = 0.94).

In addition to the just-experienced transitions, we also separately analysed the replay of all other transitions (Fig 5H); this revealed a much broader picture, but we observed the same tendency for sub-optimal actions to be replayed more (Wilcoxon rank-sum test, *W* = 13.5, *p* ≪ 0.0001). In this case, we found that both 1-move (Wilcoxon rank-sum test, *W* = 48.9, *p* ≪ 0.0001) and 2-move (Wilcoxon rank-sum test, *W* = 12.2, *p* ≪ 0.0001) sub-optimal actions were replayed at higher action entropy values than the corresponding optimal actions. We also compared the replays of recent and ‘other’ transitions and their entropy values. We found that both optimal other (Wilcoxon rank-sum test, *W* = 16.2, *p* ≪ 0.0001) and sub-optimal other (Wilcoxon rank-sum test, *W* = 14.1, *p* ≪ 0.0001) transitions were replayed more than the recent ones. All 1-move and 2-move optimal and sub-optimal other transitions were replayed at significantly higher action entropy values than the corresponding recent transitions (1-move optimal recent vs 1-move optimal other, Wilcoxon rank-sum test, *W* = 11.9, *p* = ≪ 0.0001; 1-move sub-optimal recent vs 1-move sub-optimal other, Wilcoxon rank-sum test, *W* = 9.56, *p* ≪ 0.0001, 2-move optimal recent vs 2-move optimal other, Wilcoxon rank-sum test, *W* = 4.13, *p* = 3.63·10^−5^; 2-move sub-optimal recent vs 2-move sub-optimal other, Wilcoxon rank-sum test, *W* = 15.1, *p* ≪ 0.0001). The latter observation could potentially explain why these ‘distal’ replays of other transitions were not detectable in the study of Eldar *et al*. 2020 [31], since the content of highly entropic on-task replay events may have been improbable to identify with classifiers trained on data obtained during the pre-task stimulus exposure (while subjects were contemplating the images with certitude).

Overall, we found that the pattern of the replay choices selected by our agent with parameters fit to the data for the subject shown in Fig 5 was consistent with the observations reported by Eldar *et al*. 2020 [31]. Furthermore, this consistency was true across all subjects with significant replay (Fig 6). Each MI subject, on average, preferentially replayed sub-optimal actions (just-experienced transitions, Wilcoxon rank-sum test, *W* = 2.60, *p* = 0.009; other transitions, Wilcoxon rank-sum test, *W* = 4.16, *p* = 3.14 · 10^−5^). As for the subject in Fig 5, we assessed whether the action entropy associated with the just-experienced transition (be it optimal or sub-optimal) when it was replayed was lower than the action entropy associated with other optimal and sub-optimal actions when they were replayed. Overall, we found the same trend across all MI subjects but for all combinations: i.e., 1-move transitions (just-experienced 1-move optimal vs other 1-move optimal, Wilcoxon rank-sum test, *W* = 13.8, *p* ≪ 0.0001; just-experienced 1-move sub-optimal vs other 1-move sub-optimal, Wilcoxon rank-sum test, *W* = 18.0, *p* ≪ 0.0001) and 2-move transitions (just-experienced 2-move optimal vs other 2-move optimal, Wilcoxon rank-sum test, *W* = 27.9, *p* ≪ 0.0001; just-experienced 2-move sub-optimal vs other 2-move sub-optimal, Wilcoxon rank-sum test, *W* = 53.7, *p* ≪ 0.0001). Furthermore, our agent predicted that, in each trial, MI human subjects increased their (objective) average value function by 0.196 reward points (1-sample 2-tailed t-test, *t* = 2.79, *p* = 0.011, Fig 6C) as a result of replay. As a final measure of the replay benefit, we quantified the average trial-wise increase in the probability of choosing an optimal action as a result of replay (Fig 6D). This was done to provide a direct measure of the effect of replay on the subjects’ policy, as opposed to the proxy reported by Eldar *et al*. 2020 [31] in terms of the visitation frequency (shown in Fig 1B). On average, our modelling showed that replay increased the probability of choosing an optimal action in MI subjects by 3.36% (1-sample 2-tailed t-test, *t* = 3.36, *p* = 0.003), which is very similar to the numbers reported by Eldar *et al*. 2020 [31].

**Fig. 6.**
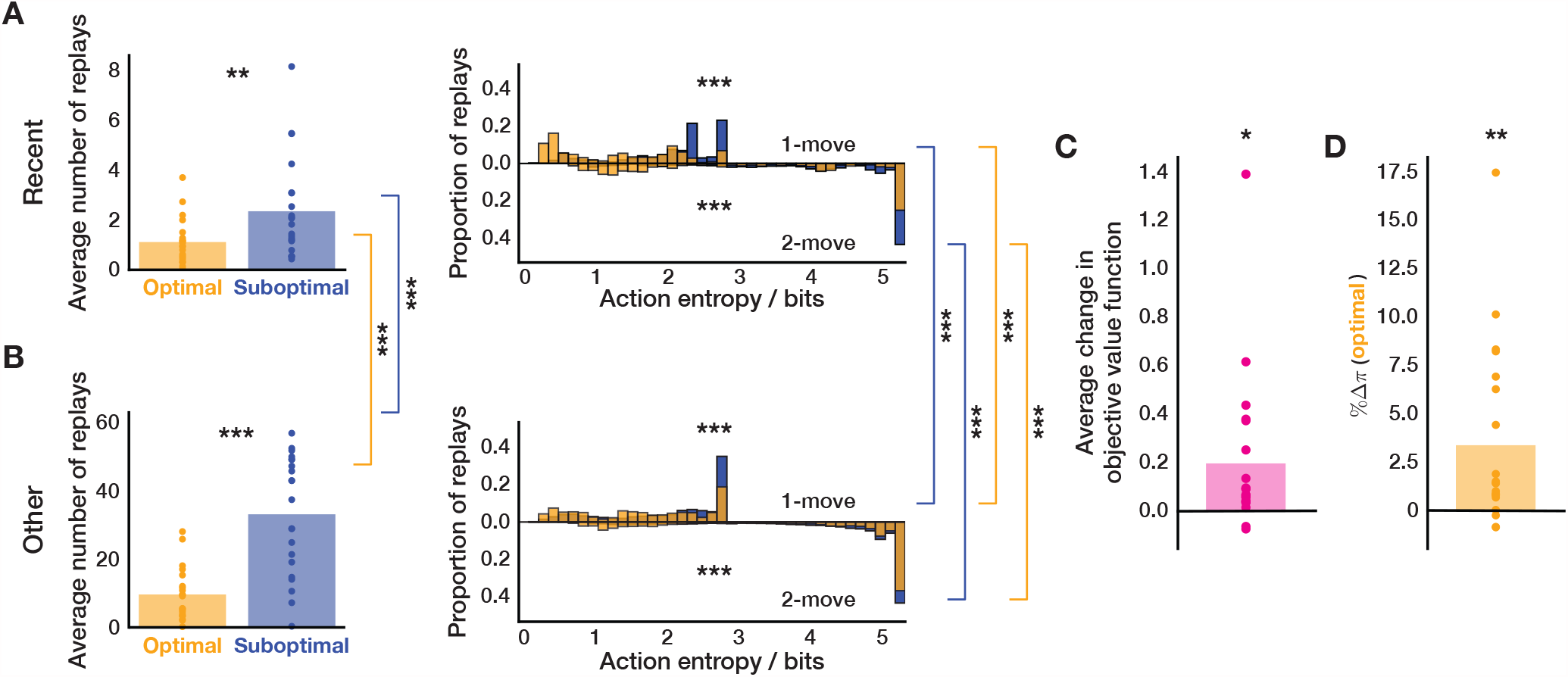
Overall On-Task Replay Statistics Across MI Subjects. (A) Left: average number of replays of just-experienced optimal and sub-optimal actions; right: proportion of action entropy values at which just-experienced optimal and sub-optimal actions were replayed. (B) Same as above but for other transitions. (C) Average change in objective value function due to replay. (D) Average change in the probability of selecting an (objectively) optimal action due to replay. * *p <* 0.05, ** *p <* 0.01, *** *p <* 0.001.

Clearly, our modelling predicted quite a diverse extent to which MI subjects objectively benefited (or even hurt themselves) by replay (Fig 6C, D). This is because the subjects had to rely on their forgetful and imperfect state-transition models to estimate MB *Q*-values for each action, and as a result of MB forgetting, some subjects occasionally mis-estimated some sub-optimal actions to be optimal – and thus their average objective replay benefit was more modest (or even negative in the most extreme cases). Indeed, upon closer examination of our best-fitting parameter estimates for each MI subject, we noted that subjects who, on average, hurt themselves by replay (*n* = 2) had very high MB forgetting rates (S3 Fig D), such that their forgetting-degraded models further exacerbated their knowledge of the degree to which each action was rewarding (the example subject shown in S7 Fig has this characteristic). Moreover, we found a significant anticorrelation of MB forgetting parameter and IF in MI subjects (multiple regression, *β* = −0.65, *t* = −5.63, *p* ≪ 0.0001; S3 Fig F), which suggests that MI subjects with a more accurate state-transition model achieved higher decision flexibility by preferentially engaging in pessimized replay following each trial outcome.

We did not find any significant linear relationship between the gain threshold and objective replay benefit (multiple regression, *β* = −0.04, *t* = −0.38, *p* = 0.710; S3 Fig F). This suggests that the effect is either non-linear or highly dependent on the current state of knowledge of the agent (due to the on-policy nature of replay in our algorithm), and thus higher MB forgetting rate could not have been ameliorated by simply having a stricter gain threshold. By contrast, we observed that the gain threshold correlated significantly with IF (multiple regression, *β* = 0.64, *t* = 4.42, *p* = 0.001; S3 Fig B). Therefore, we confirm that the gain threshold plays an important role – if it is too low, then the MB system will train the MF policy extensively, which will be problematic if the model is subject to substantial forgetting.

The apparent exuberance of distal replays that we discovered suggested that our model fit predicted the subjects to be notably more forgetful than found by Eldar *et al*. 2020 [31]. We thus compared our best-fitting MB and MF forgetting parameter estimates to those reported by Eldar *et al*. 2020 [31] for the best-fitting model. We found that our model predicted most subjects to be subject to far less MB forgetting (S2 Fig B). On the other hand, our MF forgetting parameter estimates implied that the subjects remembered their MF *Q*-values significantly less well (Supplementary Fig 2A). We (partially) attribute these differences to optimal replay: in Eldar *et al*. 2020 [31] the state-transition model explicitly affected every choice, whereas in our model, the MF (decision) policy was only informed by MB quantities so long as this influence was estimated to be gainful. Thus, the MF policy could be more forgetful, and yet be corrected by subjectively optimal MB information; this is also supported by the significant correlation of MF forgetting and objective replay benefit (multiple regression, *β* = 0.89, *t* = 7.76, *p* ≪ 0.0001; S3 Fig F). Moreover, due to substantially larger MF forgetting predicted by our modelling, we predicted that the subjects would have been more ‘surprised’ by how rewarding each outcome was. The combination of this surprise with the (albeit reduced) suprise about the transition probabilities, arising from the (lower) MB forgetting, resulted in surplus distal replays.

## Discussion

In summary, we studied the consequences of forgetting in a DYNA-like agent [39] with optimised replay [41]. The agent uses on-line experiences to train both its model-free policy (by learning about the rewards associated with each action) and its model-based system (by (re)learning the transition probabilities). It uses off-line experience to allow MF values to be trained by the MB system in a supervised manner. Behaviour is ultimately controlled exclusively by the MF system. The progressive inaccuracy of MF values can be ameliorated by MB replay, but only if the MB system has itself not become too inaccurate.

In particular, we showed that the structure of forgetting could favour the replay of sub-optimal rather than optimal actions. This arose through the interaction of several factors. One is that MF values relax to the mean experienced reward (meaning that sub-optimal actions come to look better, and optimal actions look worse than they actually are) – this can lead to the progressive choice of sub-optimal actions, and thus gain in suppressing them. A second is that optimal actions generally enjoy greater updating from actual on-line experience, since they are chosen more frequently. A third is that MB values relax towards the mean of all the rewards in the environment (because the transition probabilities relax towards uniformity), which is pessimistic relative to the experienced rewards. This makes the model relatively worse at elevating optimal actions in the MF system. We showed that the Shannon entropy of the transition distribution was a useful indicator of the status of forgetting.

We found that replay can both help and hurt, from an objective perspective – with the latter occurring when MB forgetting is too severe. This shows that replay can be dangerous when subjects lack meta-cognitive monitoring insight to be able to question the veracity of their model and thus the benefit of using it (via subjective gain estimates). This result has implications for sub-optimal replay as a potential computational marker of mental dysfunction [51, 52].

We studied these phenomena in the task of Eldar *et al*. 2020 [31], where it was initially identified. Forgetting had already been identified by Eldar *et al*. 2020 [31]; however, it played a more significant role in our model, because we closed the circle of having MB replay affect MF values and thus behaviour. By not doing this, forgetting may have been artificially downplayed in the original report (S2Fig A, B). We fit the free parameters of our agent to the behavioural data from individual subjects, correctly capturing their behavioural flexibility (IF) (S3 Fig A, B), and providing a mechanistic explanation for the replay choice preferences of those employing a hybrid MB/MF strategy. Our fits suggested that these MI subjects underwent significantly more MF forgetting than those whose behavior was more purely MF (since the latter could only rely on the MF system, see S3 Fig C).

Our study has some limitations. First, we shared Mattar and Daw 2018’s [41] assumption that the subjects could compute gain correctly in their resulting incorrect models of the task. How gain might actually be estimated and whether there might be systematic errors in the calculations involved are not clear. However, we note that spiking neural networks have recently proven particularly promising for the study of biologically plausible mechanisms that support inferential planning [53, 54], and they could therefore provide insights into the underlying computations that prioritise the replay of certain experiences.

Second, our model did not account for the need term [41] that was theorised to be another crucial factor for the replay choices. This was unnecessary for the current task. Need is closely related to the successor representation (SR) of sequential transitions [49] inasmuch as it predicts how often one expects each state to be visited given the current policy. Need was shown to mostly influence forward replay at the very outset of a trial as part of planning, something that is only just starting to be detectable using MEG [55].

The SR also has other uses in the context of planning, and is an important intermediate representation for various RL tasks. Furthermore, it has a similar computational structure to MF *Q*-values – in fact, it can be acquired through a form of MF Q learning with a particular collection of reward functions [49]. Thus, it is an intriguing possibility, suggested, for instance, in the SR-DYNA algorithm of Russek *et al*. 2017 [56] or by the provocative experiment of Carey *et al*. 2019 [57] that there may be forms of MB replay that are directed at maintaining the integrity and fidelity of the SR in the face of forgetting and environmental change. Indeed, grid cells in the superficial layers of entorhinal cortex were shown to engage in replay independently of the hippocampus, and thus could be a potential candidate for SR-only replay [58, 59]. More generally, other forms of off-line consolidation could be involved in tuning and nurturing cognitive maps of the environment, leading to spatially-coherent replay [60].

Third, Eldar *et al*. 2020 [31] also looked at replay during the 2-minute rest period that preceded each block, finding an anticorrelation between IF and replay. The most interesting such periods preceded blocks 3 and 5, after the changes to the reward and transition models. Before block 3, MF subjects replayed transitions that they had experienced in block 2 and preplayed transitions they were about to choose in block 3; MI subjects, by contrast, showed no such bias. Although we did show that even if we set the MF *Q*-values to zero before allowing the agent to engage in replay prior to starting blocks 3 and 5 (S7 Fig), we still recover our main conclusions, the fact that we do not know what sort of replay might be happening during retraining meant that we left modelling this rest-based replay to future work.

To summarize, in this work, we showed both in simulations and by fitting human data in a simple planning task that pessimized replay can have distinctly beneficial effects. We also showed the delicacy of the balance of and interaction between MB and MF systems when they are forgetful – something that will be of particular importance in more sophisticated and non-deterministic environments that involve partial observability and large state spaces [61, 62].

## Materials and Methods

### Task Design

We simulated the agent in the same task environment and trial and reward structure as Eldar *et al*. 2020 [31]. The task space contained 8 states arranged in a 2 × 4 torus, with each state associated with a certain number of reward points. From each state, 4 actions were available to the agent – up, down, left, and right. If the state the agent transitioned to was revealed (trials with feedback), the agent was awarded reward points associated with the state to which it transitioned.

The agent started each trial in the same location as the corresponding human subject, which was originally determined in a pseudo-random fashion. The agent then chose among the available moves. Based on the chosen move, the agent transitioned to a new state and received the reward associated with that state. In 1-move trials, this reward signified the end of the trial, whereas in 2-move trials the agent then had to choose a second action either with or without feedback as to which state the first action had led. In addition, transitions back to the start state in 2-move trials were not allowed. Importantly, the design of the task ensured that optimal first moves in 2-move trials were usually different from optimal moves in 1-move trials.

As for each of the human subjects, the simulation consisted of 5 task blocks that were preceded by 6 training blocks. Each training block consisted of twelve 1-move trials from 1 of 2 possible locations. The final training block contained 48 trials where the agent started in any of the 8 possible locations. In the main task, each block comprised 3 epochs, each containing six 1-move trials followed by twelve 2-move trials, therefore giving in total 54 trials per block. Every 6 consecutive trials the agent started in a different location except for the first 24 2-move trials of the first block, in which Eldar *et al*. 2020 [31] repeated each starting location for two consecutive trials to promote learning of coupled moves. Beginning with block 2 the agent did not receive any reward or transition feedback for the first 12 trials of each block. After block 2 the agent was instructed about changes in the reward associated with each state. Similarly, before starting block 5 the agent was informed about a rearrangement of the states in the torus. Eldar *et al*. 2020 [31] specified this such that the optimal first move in 15 out of consecutive 16 trials became different.

### DYNA-like Agent

#### Free Parameters

Our model has the following potential free parameters:

|  |  |
| --- | --- |
| $\eta^{MF1}$ | learning rate in 1-move trials and second moves in 2-move trials |
| $\eta^{MF2}$ | learning rate in first moves in 2-move trials |
| $\eta_r^{MF1}$ | learning rate for replay in 1-move trials and second moves in 2-move trials |
| $\eta_r^{MF2}$ | learning rate for replay in first moves in 2-move trials |
| $\beta_1^{MF1}$ | inverse temperature in 1-move trials |
| $\beta_2^{MF1}$ | inverse temperature for second moves in 2-move trials |
| $\beta_2^{MF2}$ | inverse temperature for first moves in 2-move trials |
| $\theta$ | initialisation mean for $Q^{MF}$ values |
| $\gamma_m$ | fixed bias for each action, subject to $\sum_m \gamma_a = 0$ |
| $\eta^{MB}$ | state-transition model learning rate |
| $\rho$ | fraction of the state-transition model learning rate that is used for updating opposite transitions |
| $\phi^{MB}$ | state-transition model forgetting |
| $\phi'^{MB}$ | state-transition model forgetting upon spatial re-arrangement |
| $\omega$ | state-transition model re-arrangement success |
| $\phi^{MF}$ | $Q^{MF}$ values forgetting |
| $\phi'^{MF}$ | $Q^{MF}$ values forgetting upon spatial re-arrangement |
| $\xi$ | gain threshold for initiating replay |

#### Choices

The agent has the same model-free mechanism for choice as designed by Eldar *et al*. 2020 [31]. However, unlike Eldar *et al*. 2020 [31], choices are determined exclusively by the MF system; the only role the MB system plays is via replay, updating the MF values.

The model-free system involves two sets of state-action or *Q*^*MF*^ values [48]: *Q*^*MF*1^ for single moves (the only move in 1-move trials, and both first and second moves in 2-move trials), and *Q*^*MF*2^ for coupled moves (2-move sequences) in 2-move trials.

In 1-move trials, starting from state *s*_*t*,1_, the agent chooses actions according to the softmax policy:

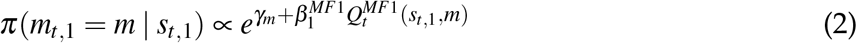

where *m ∈* {up, down, left, right}. In 2-move trials, the first move is chosen based on the combination of both sorts of *Q*^*MF*^ values:

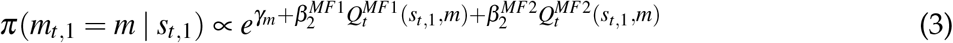

where the individual *Q*^*MF*1^ value is weighted by a different inverse temperature or strength parameter 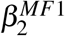, and the coupled move 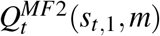 value comes from considering all possible second moves *m*_*t*,2_, weighted by their probabilities:

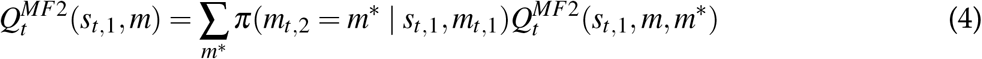

When choosing a second move the agent takes into partial account (since *Q*^*MF*2^ doesn’t) the state *s*_*t*,2_ to which it transitioned on the first move:

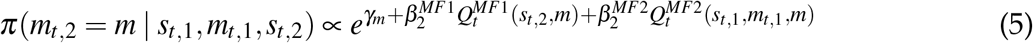

In trials without feedback, since the agent’s *Q*^*MF*^ values are indexed by state and the transition is not revealed, it is necessary in equation 5 to average *Q*^*MF*1^ over all the permitted *s*_*t*,2_ (since returning to *s*_*t*,1_ is disallowed):

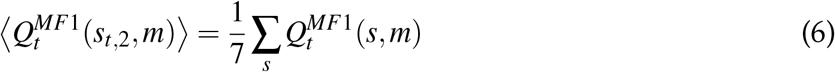

The decision is then made according to equation 5.

#### Model-Free Value Learning

The reward *R*(*s*_*t*,2_) received for a given move *m*_*t*_ and transition to state *s*_*t*,2_ is used to update the agent’s *Q*^*MF*1^ value for that move:

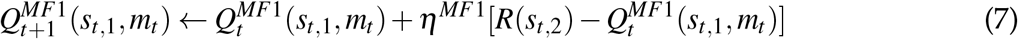

On 2-move trials, the same rule is applied for the second move, adjusting 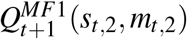 according to the second reward *R*(*s*_*t*,3_). Furthermore, on 2-move trials, *Q*^*MF*2^ values are updated at the end of each trial based on the sum total reward obtained on that trial but with a different learning rate:

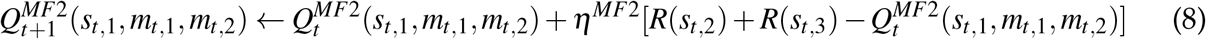

#### Model-Based Learning

Additionally, and also as in Eldar *et al*. 2020 [31], the agent’s model-based system learns about the transitions associated with every move it experiences (i.e. the only transition in 1-move trials and both transitions in 2-move trials):

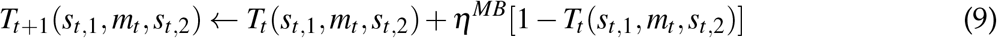

and, given that actions are reversible, learning also happens for the opposite transitions to a degree that is controlled by parameter *ρ*:

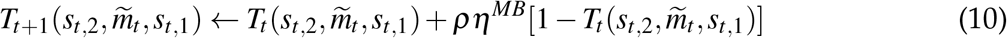

where 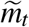 is the transition opposite to *m*_*t*_. Note that in trials with no feedback the agent does not receive any reward and the state it transitions to is uncued. Therefore, no learning occurs in such trials.

To ensure that the probabilities sum up to 1, the agent re-normalizes the state-transition model after every update and following the MB forgetting (see below) as:

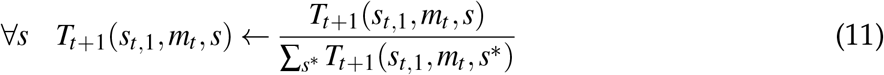

#### Replay

That our agent exploits replay is the critical difference from the model that Eldar *et al*. 2020 [31] used to characterize their subjects’ decision processes.

Our algorithm makes use of its (imperfect) knowledge of the transition structure of the environment to perform additional learning in the inter-trial intervals by means of generative replay. Specifically, the agent utilises its state-transition model *T* and reward function *R*(*s*) to estimate model-based 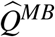 values for every possible action (that are allowable according to the model). These 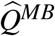 values are then assessed for the potential MF policy improvements (see below).

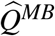 values for 1-move trials and second moves in 2-move trials are estimated as follows:

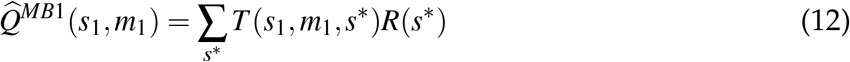

Similarly, 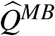 values for 2-move sequences are estimated as:

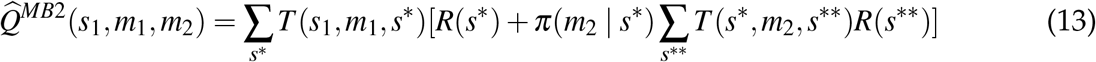

where *π*(· |*s*) is the agent’s MF policy at state *s* (computed according to equation 2) which has been updated by on-line learning in the immediately preceding trial. When summing over the potential outcomes for a second action in equation 13, the agent additionally sets the probability of transitioning into the starting location *s*_1_ to zero (since back-tracking was not allowed) and normalises the transition probabilities according to equation 11. Note that reward function *R*(*s*) here is the true reward the agent would have received for transitioning into state *s* since we assume that the subjects have learnt the image-reward associations perfectly well. The model-generated 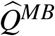 values therefore incorporate the agent’s uncertainty about the transition structure of the environment. If the agent is certain which state a given action would take it to, 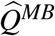 value for that action would closely match the true reward function of that state. Otherwise,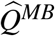 values for uncertain transitions are corrupted by the possibility of ending up in different states.

The agent then uses all the generated 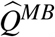 values to compute new hybrid *Q*^*MF/MB*^ These hybrid values correspond to the values that would have resulted had the current *Q*^*MF*^ values. values been updated towards the model-generated 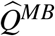 values using replay-specific learning rates 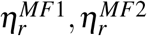:

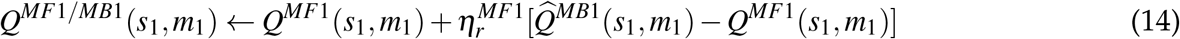

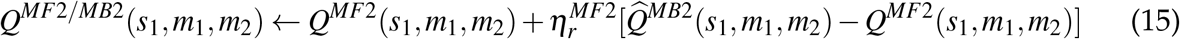

We note that the 2-move updates specified in equation 15 are a form of supervised learning, and so differ from the RL/DYNA-based episodic replay suggested by Mattar and Daw 2018 [41]. As mentioned above, we chose this way of updating MF *Q*-values for coupled moves in replay to keep the algorithmic details as close to Eldar *et al*. 2020 [31] as possible. In principle, we could have also operationalised our 2-move replay in a DYNA fashion.

To assess whether any of the above updates improve the agent’s MF policy, the agent computes the expected value of every potential update [41]:

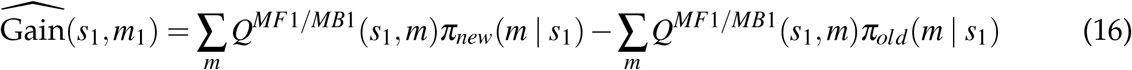

Analogously, for a sequence of two moves *{m*_1_, *m*_2_*}*:

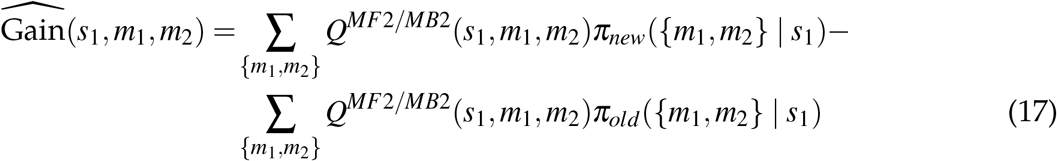

where the policy *π* was assumed to be unbiased and computed as in equation 2; that is, the agent directly estimated the corresponding probabilities for each sequence of 2 actions in 2-move trials.

Both of these expressions for estimating 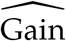 use the full *new* model-free policy that would be implied by the update. Thus, as also in Mattar and Daw 2018 [41], the gain (equations 16 and 17) does not assess a psychologically-credible gain, since the new policy is only available *after* the replays are executed. Moreover, we emphasise that this same gain is the agent’s estimate, for the true gain is only accessible to an agent with perfect knowledge of the transition structure of the environment (which is infeasible in the presence of substantial forgetting).

Finally, the expected value of each backup (EVB), or replay, is computed as 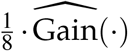 (since the agent starts each trial in a pseudorandom location, we assumed need to be uniform). Exactly as in Mattar and Daw 2018 [41], the priority of the potential updates is determined by the EVB value – if the greatest EVB value exceeds gain threshold *ξ* (for simplicity and due to the assumption of uniform need, we refer to *ξ* as gain threshold, rather than EVB threshold), then the agent executes the replay associated with that EVB towards the model-generated 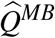 value according to equations 14 or 15 (depending whether it is a single move or a two-move sequence), thus incorporating its MB knowledge into the current MF policy. Note that since this changes the agent’s MF policy and the generation of 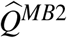 is policy-dependent, the latter are re-generated following every executed backup. The replay proceeds until no potential updates have the EVB value greater than *ξ*.

#### Forgetting

The agent is assumed to forget both about the *Q*^*MF*^ values and the state-transition model *T* in trials where feedback is provided. Thus, after every update and following replay the agent forgets according to:

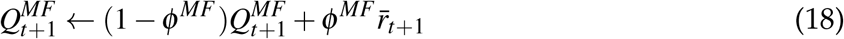

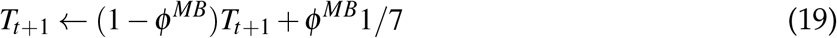

Note that we parameterize the above equations in terms of forgetting parameters *ϕ*. Eldar *et al*. 2020 [31] instead used *τ* as remembrance, or value retention, parameters. Therefore, our forgetting parameters *ϕ* are equivalent to Eldar *et al*. 2020 [31]’s 1 − *τ*.

The state-transition model therefore decays towards the uniform distribution over the potential states the agent can transition to given any pair of state and action. *Q*^*MF*^ values are forgotten towards the average reward experienced since the beginning of the task, 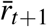. For *Q*^*MF*1^ values, it is the average reward obtained in single moves:

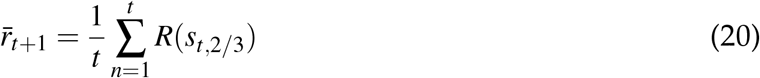

and for *Q*^*MF*2^ values it is the average reward obtained in coupled moves:

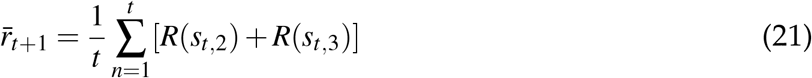

where *R*(*s*) is the reward obtained for transitioning into state *s*. This differs from Eldar *et al*. 2020 [31], for which forgetting was to a constant value *θ*, which was a parameter of the model.

After blocks 2 and 4 the learnt *Q*^*MF*^ values are of little use due to the introduced changes to the environment, and, again as in Eldar *et al*. 2020 [31] the agent forgets both the *Q*^*MF*^ values and state-transition model *T* according to equations 18 and 19, but with different parameters *ϕ* ′^*MF*^ and *ϕ* ′^*MB*^, respectively.

In addition to on-task replay, the agent also engages in off-task replay immediately before the blocks with changed image-reward associations and spatial re-arrangement. In the former case, the agent uses the new reward function *R*′ (*s*) that corresponds to the new image-reward associations. In the latter case, the agent generates 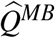 values with the new reward function and a state-transition model rearranged according to the instructions albeit with limited success:

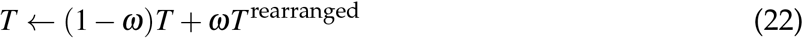

During these two off-task replay bouts, the agent uses the exact same subject-specific parameter values as in on-task replay. The effect of such model re-arrangement on the accuracy of 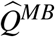 estimates of the re-arranged environment in MI human subjects, as modelled by our agent, is shown in S6 Fig.

#### Initialisations

*Q*^*MF*^ values are initialised to *Q*^*MF*^ ∼ 𝒩 (*θ*, 1), and the state-transition model *T* (*s*_*t*,1_, *m*_*t*_, *s*_*t*,2_) is initialised to 1/7 (since self-transitions are not allowed). The agent, however, starts the main task with extensive training on the same training trials that the subjects underwent before entering the main task. In S8 Fig, we show how ignorant (according to our agent’s prediction) each subject was as regards the optimal transitions in the state-space after this extensive training.

### Parameter Fitting

To fit the aforementioned free parameters to the subjects’ behavioural data we used a slightly altered Markov Chain Monte Carlo sampling in the Approximate Bayesian Computation [63] framework (MCMC ABC). As a distance (negative log-likelihood) measure for ABC, we took the root-mean-squared deviation between the simulated and the subjects’ performance data, measured as the proportion of available reward collected in each epoch. The pseudocode for our fitting procedure is provided in algorithm 1.

**Algorithm 1.**
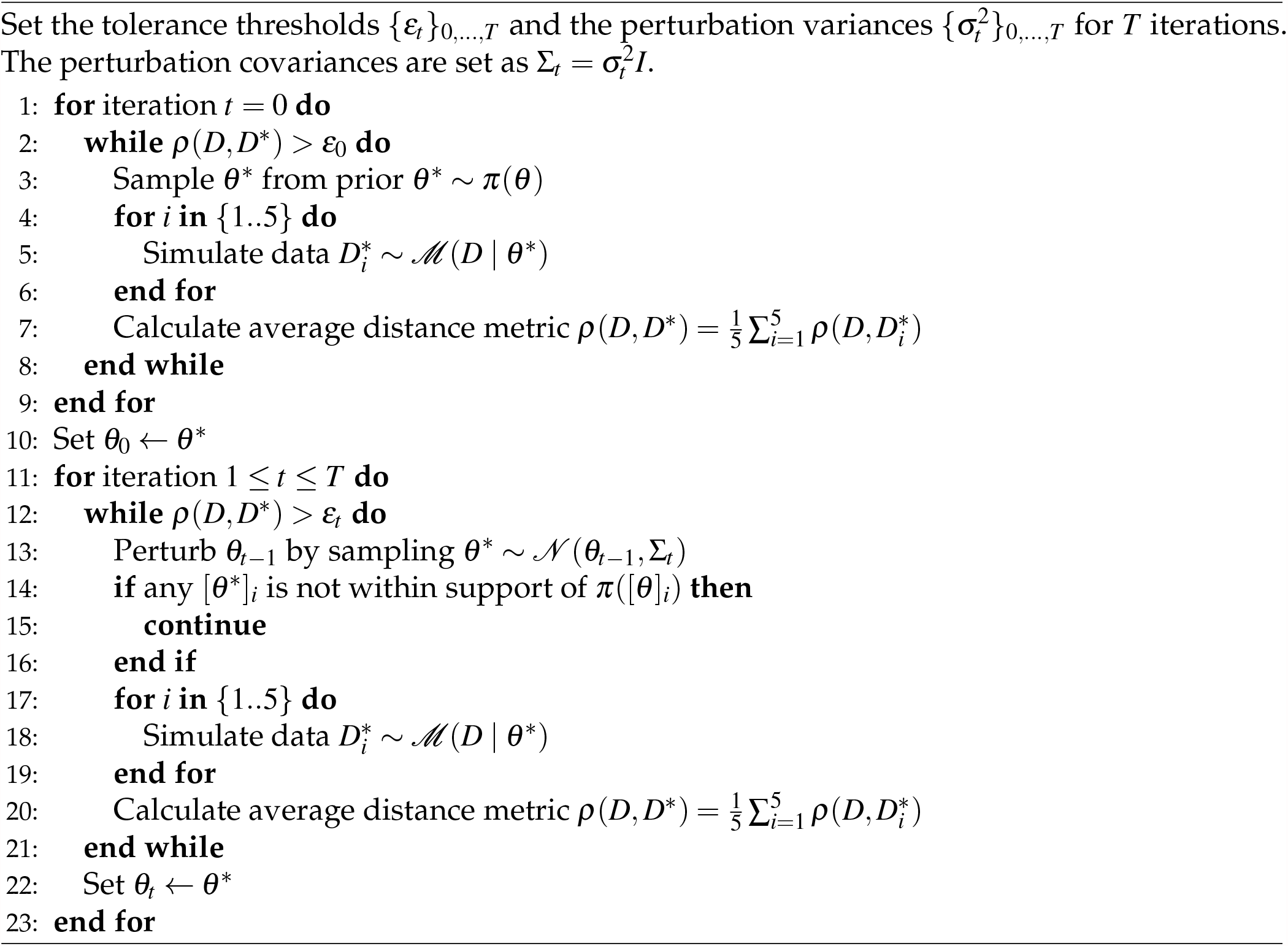
MCMC ABC Algorithm for obtaining a point estimate from a mode of the posterior distribution in the parameter space given the prior distribution *π*(*θ*), data *D*, and a model that simulates the data ℳ (*D* | *θ*). *θ*_*t*_ denotes the full multivariate parameter sample at iteration *t*.

The fitting procedure was performed for 55 iterations with the exponentially decreasing tolerance threshold *ε*_*t*_ ranging between 0.6 and 0.10. For covariance matrices, we used identity multiplied by a scalar variance with the exponential range between 0.5 and 0.01. To avoid spurious parameter samples being accepted, we simulated our model 5 times with each proposed parameter sample and then used the average over these simulations to compute the distance metric.

We chose uniform priors with support between 0 and 1 for 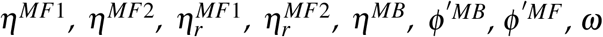, The priors for on-line forgetting parameters *ϕ* ^*MF*^ and *ϕ* ^*MB*^ were specified to be beta with *α* = 6 and *β* = 2. Parameters *θ* and *γ*_*m*_ were chosen to be Gaussian with mean 0 and variance 1. All the other parameters had gamma priors with location 1 and scale 1, except for *ξ*. Since our perturbation covariance matrices were identities multiplied by a scalar variance, and the values for *ξ* were in general very small compared to the other parameters – to allow for a similar perturbation scale for *ξ* we sampled it from a log-gamma distribution with location −1 and scale 0.01. Our fitting algorithm therefore learnt log_10_ *ξ* rather than *ξ* directly. Additionally, parameters with bounded support (such as the learning and forgetting rates) were constrained to remain within the support specified by their corresponding prior distributions.

The fitting procedure was fully parallelized thanks to the implemented python MPI module in a freely available python package *astroabc* [64]. The distribution of the resulting fitting errors is shown in S9 Fig.

### Replay Analysis

The objective replay benefit for Fig 5E was computed as the total accrued gain for all example 1-move replay events shown in Fig 5D. That is, we used equation 16, where *Q*^*MF/MB*^ were taken to be the true MF *Q*-values (or the true reward obtainable for each action), and *π*_*old*_ and *π*_*new*_ were the policies before and after all replay events, respectively. The average replay benefit in Fig 5F was computed in the same way as above, but the total accrued gain was averaged over all states where replay occurred (or equivalently, where there was a policy change). Importantly, in Fig 5E we only show the objective replay benefit as a result of the 1-move replay events from Fig 5D. For the overall average, in 2-move trials the 2-move replay events were also taken into account, and the total accrued gain after a 2-move trial was computed as the average over the 1-move and 2-move value function improvements.

The subjective replay benefit shown in S7 Fig E was computed in the exact same way as described above; however, for *Q*^*MF/MB*^ we took the agent’s updated MF *Q*-values after the replay events (or event) shown in S7 Fig D. The average objective value function change from S7 Fig F was computed in the same way as described above.

To analyse the agent’s preference to replay sub-optimal actions, we extracted the number of times the agent replayed sub-optimal and optimal actions at the end of each trial, which is shown in Figs 5G, H for the most recent and all other (or ‘distal’) transitions, respectively. Due to the torus-like design of the state space, some transitions led to the same outcomes (e.g. going ‘up’ or ‘down’ when at the top or bottom rows), and we therefore treated these as the same ‘experiences’. For instance, if the agent chose an optimal move ‘up’, then both ‘up’ and ‘down’ replays were counted as replays of the most recent optimal transition. In 2-move trials, a move was considered optimal if the whole sequence of moves was optimal. In such case, the replays of this 2-move sequence and the replays of the second move were counted towards the most recent optimal replays. If, however, the first move in a 2-move trial was sub-optimal, the second move could still have been optimal, and therefore both sub-optimal replays of the first move and optimal replays of the second move were considered in this case.

For all replays described above, we considered action entropy values at which these replays were executed (shown in Figs 5G, H, right column). For every 1-move replay, the corresponding action entropy was computed according to equation 1. For every 2-move replay beginning at state *s*_1_ and proceeding with a sequence of actions *m*_1_, *m*_2_, the corresponding action entropy was computed as the joint entropy:

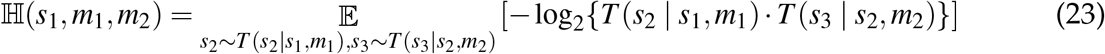

where in *T* (*s*_2_ | *s*_1_, *m*_1_) the probability of transitioning from *s*_2_ to *s*_1_ was set to zero; similarly, in *T* (*s*_3_ | *s*_2_, *m*_2_) the probabilities of transitioning from *s*_3_ to *s*_2_ and from *s*_2_ to *s*_1_ were set to zero (since back-tracking was not allowed), and the transition matrix was re-normalized as in equation 11.

### Example Simulations

The contour plot in Fig 1D was generated by simulating the agent in 1-move trials in the main state-space on a regularly spaced grid of 150 *ϕ* ^*MF*^ and *ϕ* ^*MB*^ values ranging from 0 to 0.5. For each combination of *ϕ* ^*MF*^ and *ϕ* ^*MB*^, the simulation was run 20 times for 300 trials (in each simulation the same sequence of randomly-generated starting states was used). Performance (proportion of available reward collected) in the last 100 trials of each simulation was averaged within and then across the simulations for the same combination of *ϕ* ^*MF*^ and *ϕ* ^*MB*^ to obtain a single point from the contour plot shown in Fig 1d. The final matrix was then smoothed with a 2D Gaussian kernel of std. 3.5. The model parameters used in these simulations are listed below:

|  |  |
| --- | --- |
| $\eta^{MF1}$ | 0.615 |
| $\eta_r^{MF1}$ | 0.615 |
| $\beta_1^{MF1}$ | 1.552 |
| $\eta^{MB}$ | 0.98 |
| $\rho$ | 0.70 |
| $\gamma_m$ | $[-1.105, 1.088, 0, 0.017]$ |
| $\theta$ | 0.088 |
| $\xi$ | 0.0001 |

The data from Figs 1B, 3, 4, and S1 Figs A, B were obtained by simulating the agent in a simple environment with only two actions available; in each trial the agent had to choose between the two options for 100 trials in total. The agent received a reward of 1 for the sub-optimal action and 10 for the optimal action (except for S1 Fig, there the agent was simulated with reward values that ranged from 0 to 10 with linear increments of 0.1). The following parameters were used:

|  |  |
| --- | --- |
| $\eta^{MF1}$ | 0.20 |
| $\eta_r^{MF1}$ | 0.20 |
| $\beta_1^{MF1}$ | 0.90 |
| $\eta^{MB}$ | 0.98 |
| $\phi^{MB}$ | 0.10 |
| $\phi^{MF}$ | 0.20 |
| $\xi$ | 0.00001 |

*Q*^*MF*^ values were initialised to 0, the state-transition model was initialised to 0.5, and the bias parameter *γ*_*m*_ was set to 0 for each move.

The estimated gain from Fig 3A was computed by using equation 16 where MF *Q*-values were taken to be the agent’s MF *Q*-values after the final trial, and 700 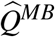 values were manually generated on the interval [−9, 12]. This procedure was repeated with the agent’s MF *Q*-values additionally decayed according to equation 18 with *ϕ* ^*MF*^ values of 0.3, 0.5 and 0.7, and the average reward obtained by the agent over those 100 trials. The x-axis was then limited to the appropriate range.

The estimated gain from Fig 4B was computed in the same way as described above, but instead of manually generating 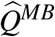 values, 100 transition probabilities (with constant linear increments) were generated on the interval from [0.5, 1] for the correct transition and [0, 0.5] for the other transition (such that the two always summed up to 1). These multiple instances of the state-transition model were used to generate 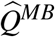 values for computing the estimated gain.

The entropy range difference in S1 Fig A was computed as the difference between the sub-optimal and optimal action entropy values at which the estimated gain became positive for each respective action after the last (100th) trial. These range differences for each combination of the reward values were then averaged over 20 simulations and plotted as a contour plot which was additionally smoothed with a 2D Gaussian kernel of 1.5 std.

The pessimism bias in S1 Fig B was computed as the average difference between the number of sub-optimal and optimal replays at the end of each trial, averaged over the same 20 simulations. The resulting matrix was also smoothed with a 2D Gaussian kernel of 1.5 std.

The contour plot in S1 Fig C was generated from the data obtained from the same simulation as for Fig 1D, and using the procedure outlined above (the bias for each combination of *ϕ* ^*MF*^ and *ϕ* ^*MB*^ values was averaged over the last 100 trials and then across 20 simulations). The resulting matrix was smoothed with a 2D Gaussian kernel of 3.5 std.

## Acknowledgements

GA, CG, and PD are funded by the Max Planck Society. PD is also funded by the Alexander von Humboldt Foundation. EE holds NIH grants R01MH124092 and R01MH125564, and a United States-Israel BSF grant 2019801.

## Competing Interests

The authors declare no competing interests.

## Supplementary Figures

**S1 Fig.**
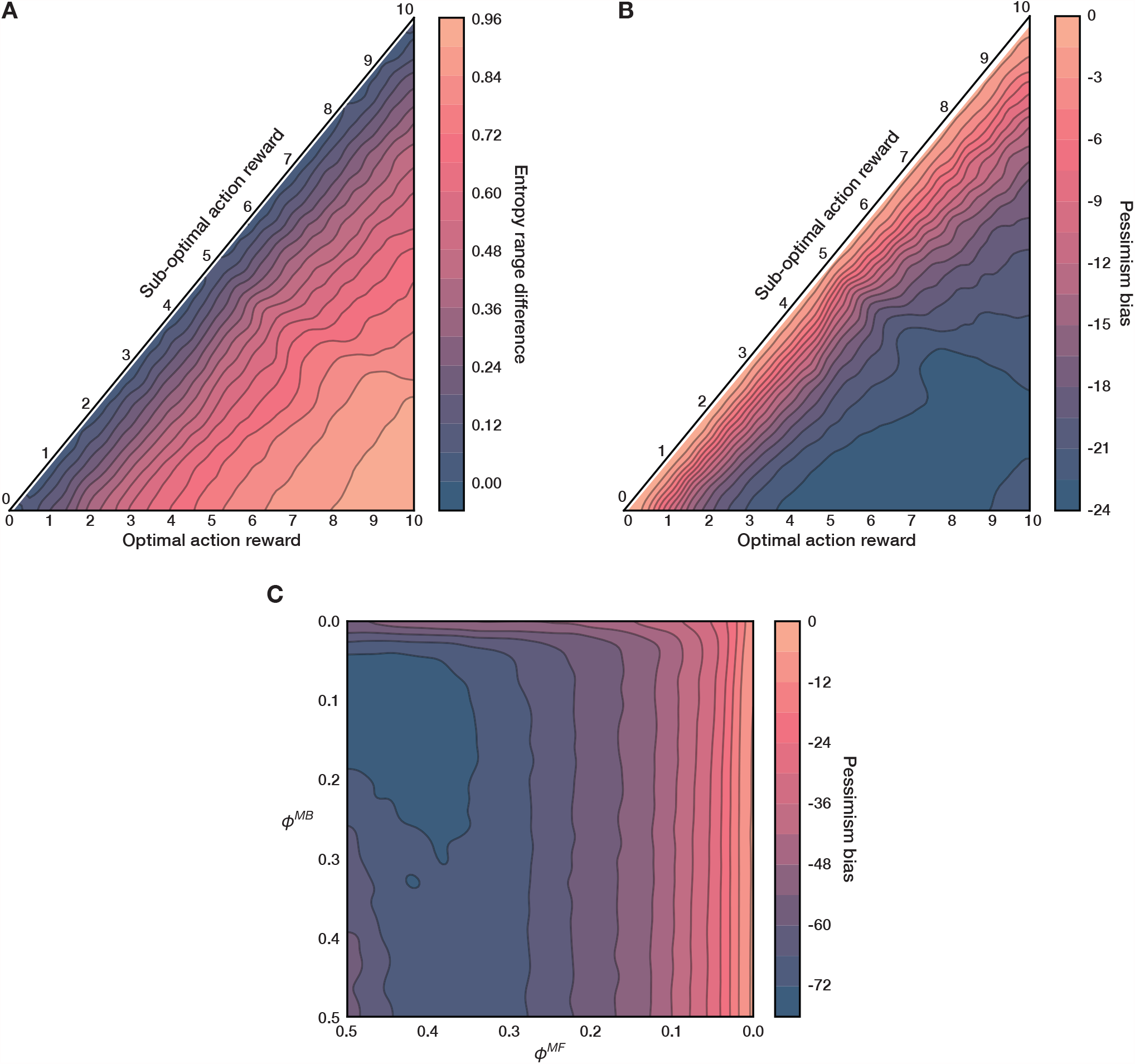
Replay Entropy Range And Pessimism Bias. (A) Difference in the range of sub-optimal and optimal action entropy values at which the estimated gain for the corresponding actions was positive. Higher values correspond to a higher action entropy range for the sub-optimal action; the agent can therefore benefit from the replay of sub-optimal actions at a wider range of action entropy values. Each datum is an average over 20 simulations where each simulation consisted of 100 trials. (B) Pessimism bias, defined here as the average (over 100 simulation trials) difference between the per-trial average number of replays of sub-optimal and optimal actions. Each datum is an average over the same 20 simulations. Negative values indicate a stronger pessimism bias. Note that both matrices are triangular, since the reward for a sub-optimal action has to be smaller than that for the optimal action (by definition); the labels on the diagonal therefore label rows. (C) Same quantity as in (B) but for the agent simulated in the behavioural task with varying MF and MB forgetting parameters.

**S2 Fig.**
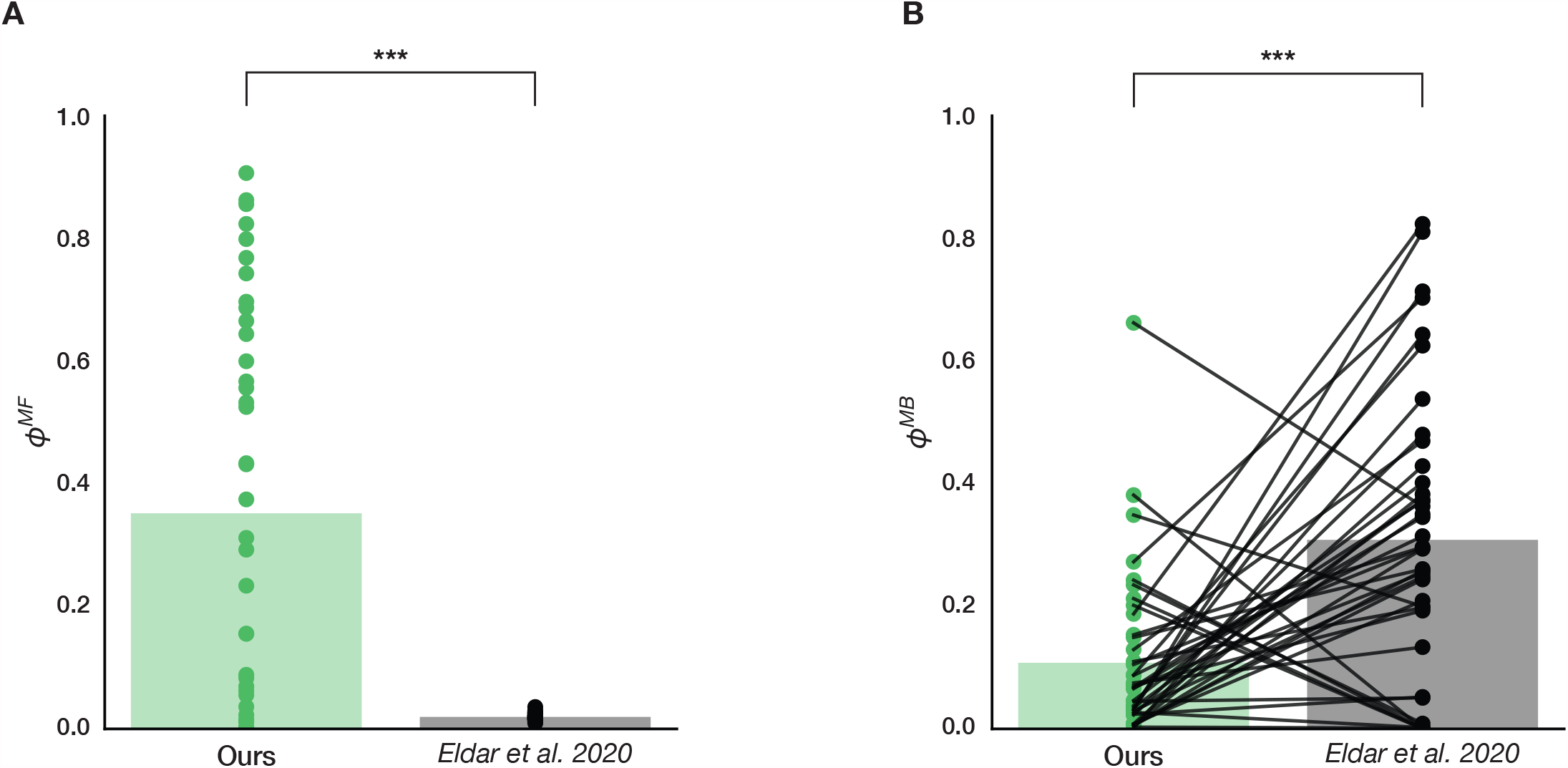
Comparison of Forgetting Parameters. (A) Comparison of our best-fitting estimates of the MF forgetting parameter, *ϕ* ^*MF*^, to that of Eldar *et al*. 2020 [31] for the hybrid MF/MB model. Our modelling predicted the subjects to be notably more forgetful as regards their MF *Q*-values (2-sample 2-tailed t test, *t* = 6.55, *p* = 5.54 · 10^−9^). (B) Same as before but for the MB forgetting parameter, *ϕ* ^*MB*^. By contrast to MF forgetting, we predicted that most of the subjects remembered their state-transition models better (2-sample 2-tailed t test, *t* = −4.76, *p* = 8.81 · 10^−6^). *** *p <* 0.0001.

**S3 Fig.**
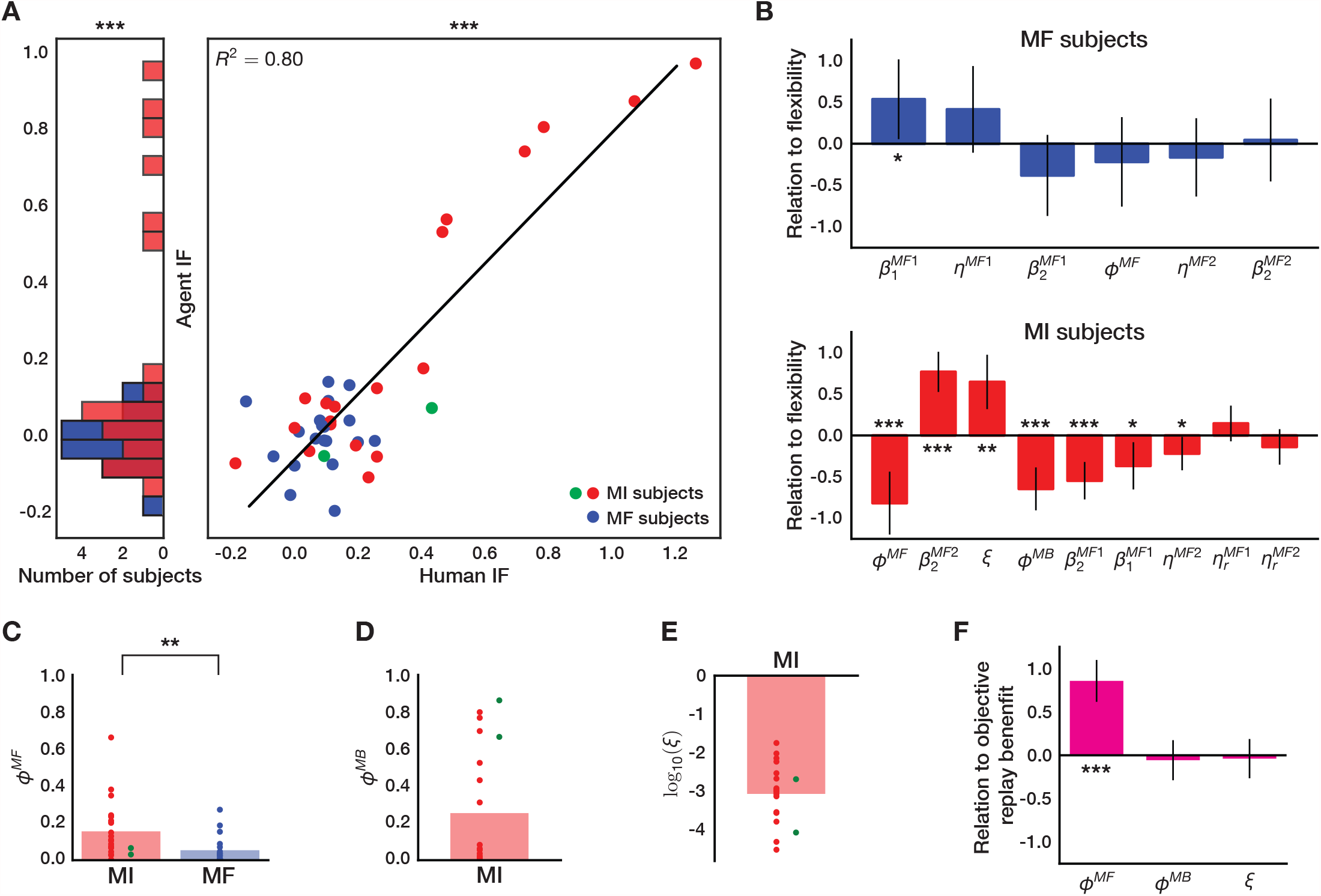
Parameter Equivalents of Subjects’ Decision Flexibility. (A) Linear regression of behavioural individual flexibility (IF) for all subjects, as measured by Eldar *et al*. 2020 [31], against IF (averaged over 100 simulations) computed from our agent with subject-specific parameters (Pearson’s coefficient of determination, *R*^2^ = 0.80, *p* ≪ 0.0001). Subjects for which our model predicted sufficient replay (MI subjects) are highlighted in red, and MF subjects are shown in blue. Further, MI subjects who were found to mostly hurt themselves by replay (*n* = 2) are shown in green. IF of MI subjects, as predicted by our agent, was significantly higher than that of MF subjects (permutation test, 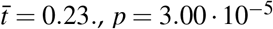, *p* = 3.00 · 10^−5^). (B) Multiple regression of all subjects’ best-fitting decision, online learning and memory parameters against their respective IF values as predicted by our agent (due to our fitting procedure, we excluded all MB-relevant parameters for MF subjects that did not engage in replay). Vertical black lines show 95% confidence intervals. (C) MF forgetting parameter of MI subjects was found to be significantly higher that that of MF subjects (Wilcoxon rank-sum test, *W* = 2.96, *p* = 3.02 · 10^−3^). (D) Distribution of MB forgetting parameter, *ϕ* ^*MB*^, in MI subjects. (E) Best-fitting gain threshold statistics across MI subjects. (F) Multiple regression of parameters that controlled the amount of replay in MI subjects against the average policy change for each subject due to replay (shown in Fig 6C). * *p <* 0.05, \*\**p <* 0.01, *** *p <* 0.0001, t test (unless specified otherwise).

**S4 Fig.**
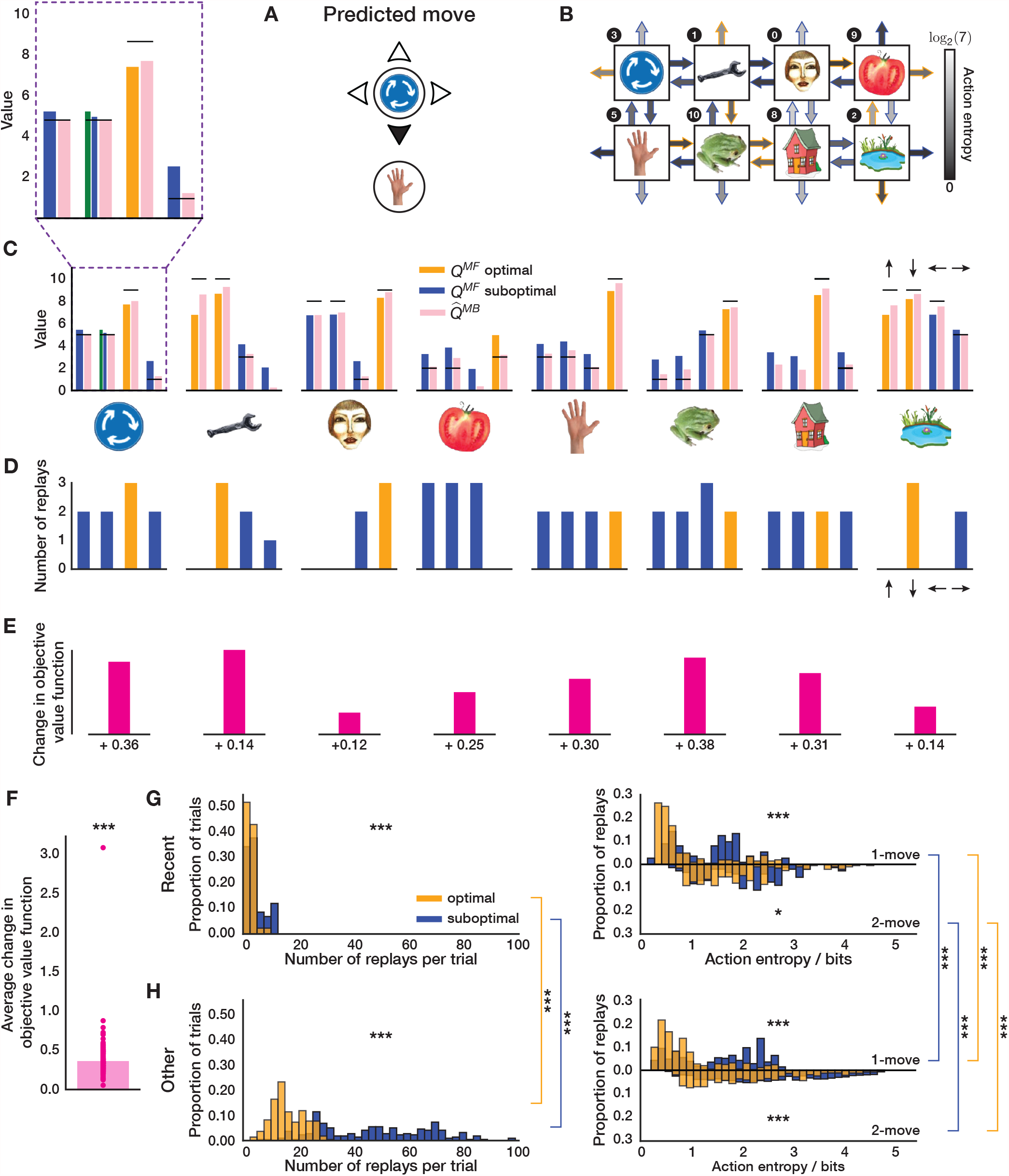
Epistemology of Replay: Another Example. The layout of the figure is similar to that of Fig 5. (A) An example move predicted by our agent with parameters fit to an MI subject. (B) State-transition knowledge of the agent after executing the move in (A). Note how the agent is less ignorant about the transitions compared to that in Fig 5. (C) MF and MB knowledge of the agent. The agent chose a sub-optimal move because its MF *Q*-value had been forgotten to an extent that made the agent more likely to choose that move. Online learning (green shading) indeed helped the agent to remember that the chosen move was worse than predicted, however only ever so slightly, for the agent’s knowledge was already fairly accurate. (D) Replays executed by the agent after learning online about the move in (A). Note that, despite the agent’s accurate MF knowledge, it still chooses to engage in replay. This is because its state-transition model was exceptionally accurate; moreover, our parameter estimates indicated that this MI subject had a very low gain threshold. (E) Changes in the objective value function as a result of the replay in (D). (F) Overall, we found that this MI subject, as predicted by our agent, achieved an average objective value function improvement of 0.37 reward points as a result of replay (1-sample 2-tailed t test, *t* = 25.2, *p* ≪ 0.0001). (G-H) Same as Fig 5. *** *p <* 0.001.

**S5 Fig.**
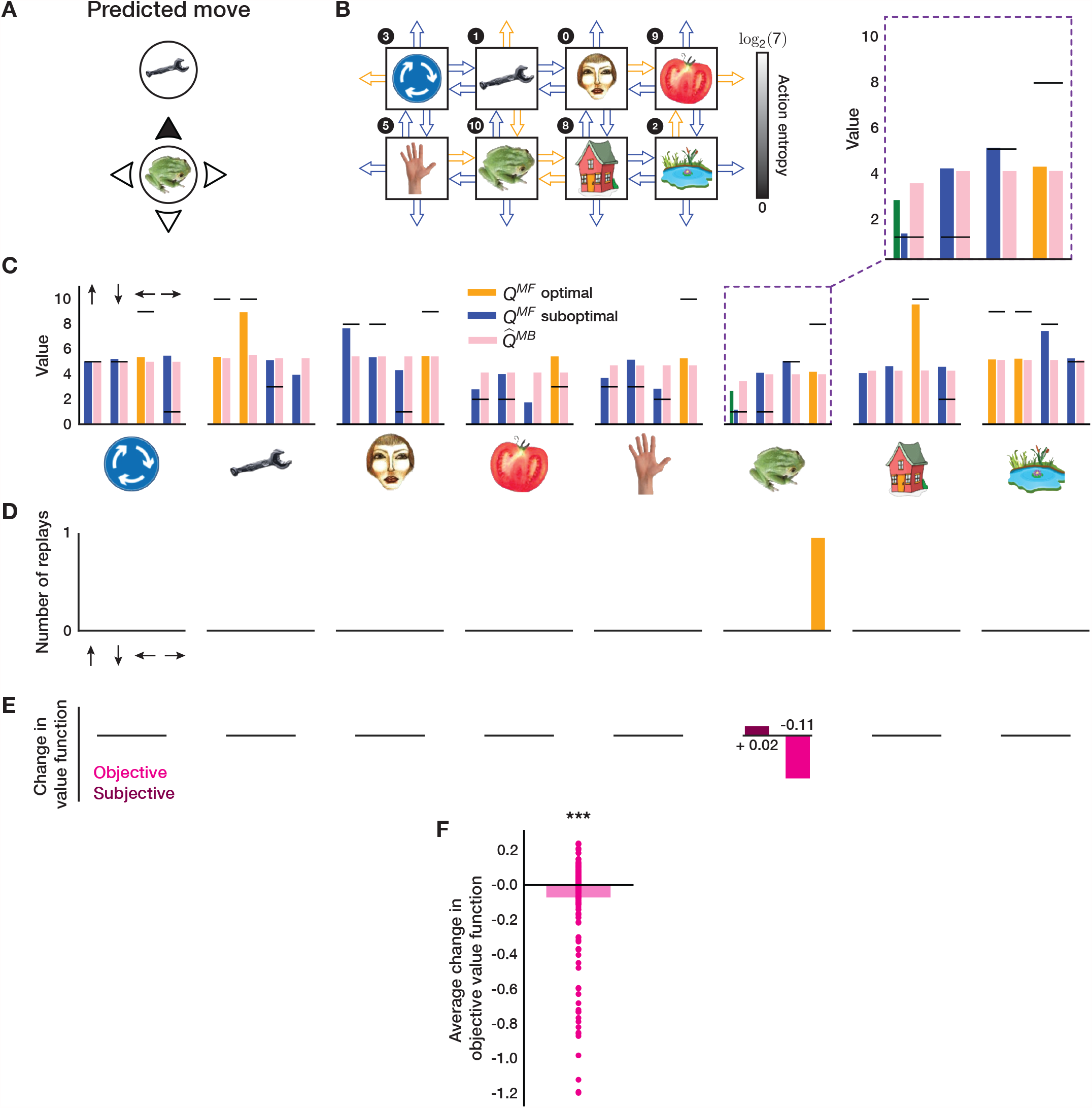
How Replay Can Hurt. The layout of the figure is similar to that of Fig 5. (A) Actual move in a 1-move trial chosen by an MI subject that was also predicted by our agent with subject-specific parameters. (B) State-transition knowledge of the agent after executing the move in (A). Note the agent’s extreme ignorance about all transitions. (C) MF and MB knowledge of the agent. The agent chose a sub-optimal move because it’s MF *Q*-values had been forgotten to an extent that made the agent more likely to choose that move. Online learning (green shading) indeed helped the agent to remember that the chosen move was worse than predicted. The agent’s MF knowledge after executing this move, however, still incorrectly indicated that the optimal move was ‘left’ (as opposed to the objectively optimal move ‘right’). (D) Replays executed by the agent after learning online about the move in (A). The agent chose to replay the move ‘right’, and such replay further exacerbated the agent’s knowledge, for it decreased the MF *Q*-value of the objectively optimal action (towards its model’s estimate, 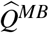, shown in pink bar). Note that although the agent replayed the objectively optimal action (‘right’) from state ‘frog’, it was in fact a subjectively sub-optimal action with respect to its state of knowledge. Such replay decreased the agent’s objective value function (as measured with respect to the true obtainable reward) of that state by 0.11 reward points (E). According to the agent’s estimate, however, it’s subjective value function of that state increased by 0.02 reward points. (F) Overall, we found that this MI subject, as predicted by our agent, mostly hurt himself by replaying at the end of each trial (average change in objective value function was decreased by 0.07 reward points, 1-sample 2-tailed t test, *t* = −4.04, *p* = 7.44 · 10^−5^). *** *p <* 0.001.

**S6 Fig.**
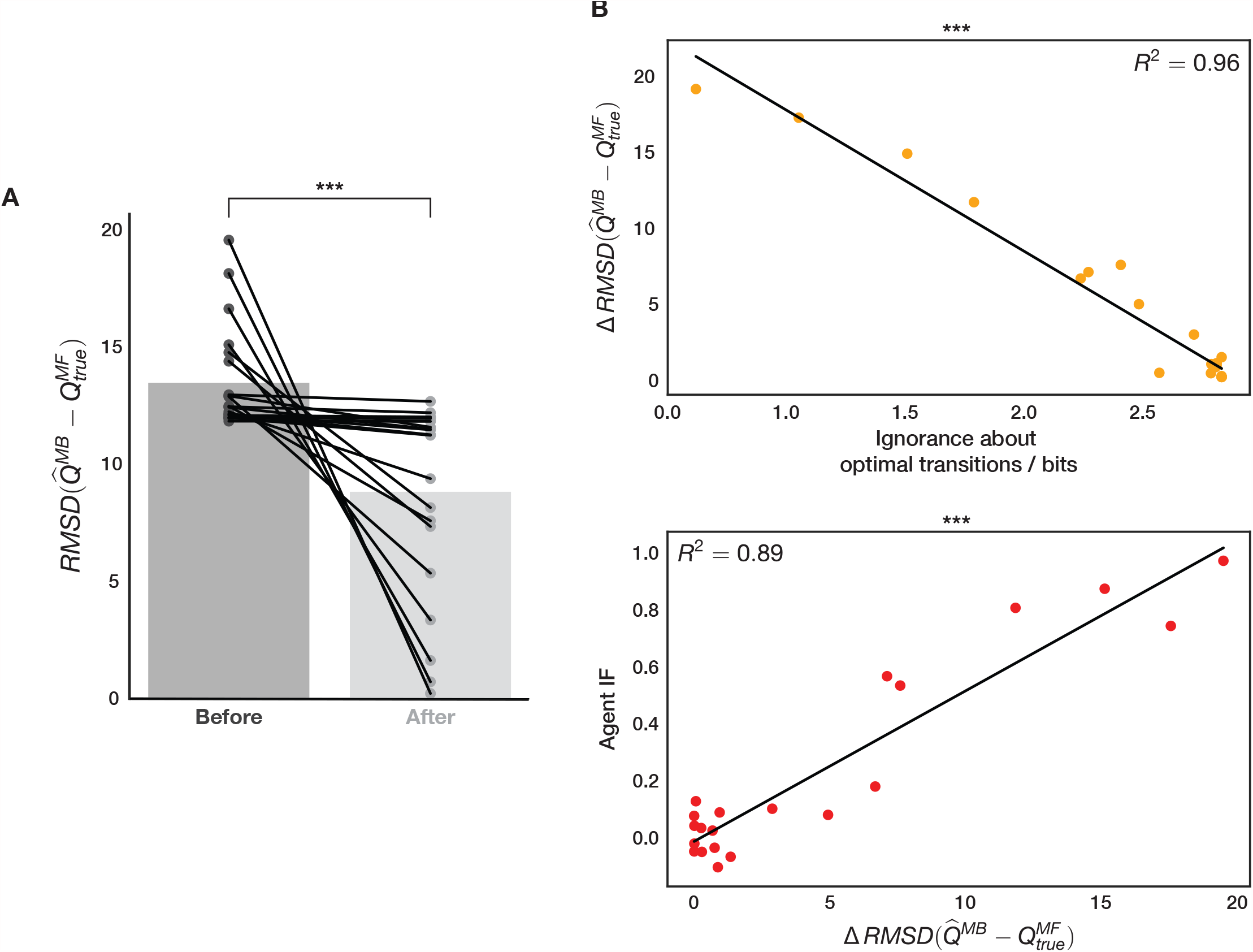
Effect of State-Transition Model Re-Arrangement. (A) Root-mean-square deviation between the MI subjects’ state-transition model (as predicted by our agent) reward estimates, 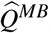, and the true reward, 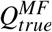, of each transition in the final block of the task before and after the model was re-arranged as in equation 22. The RMSD measure shows that the re-arrangement (even with limited success) made the state-transition model’s predictions significantly more accurate (2-sample 2-tailed t test, *t* = −4.53, *p* = 5.29 · 10^−5^). (B) Top: linear regression of the difference between the two quantities in (A) (before - after) against the ignorance about optimal transitions (measured as the average action entropy across objectively optimal actions). MI subjects with lower state-transition model entropy had significantly more accurate 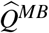 estimates as a result of the model re-arrangement (Pearson’s coefficient of determination, *R*^2^ = 0.96, *p* ≪ 0.0001.); bottom: simulated IF for MI subjects showed significant correlation to the RMSD measure (Pearson’s coefficient of determination, *R*^2^ = 0.89, *p* ≪ 0.0001). Additionally, we found that the subjects’ ignorance about objectively optimal action outcomes after the model re-arrangement correlated significantly with the simulated IF (not shown, Pearson’s coefficient of determination, *R*^2^ = 0.82, *p* ≪ 0.0001). This means that uncertainty in the subjects’ state-transition model was a good predictor of how well they adapted to the spatial re-arrangement that took place. *** *p <* 0.001.

**S7 Fig.**
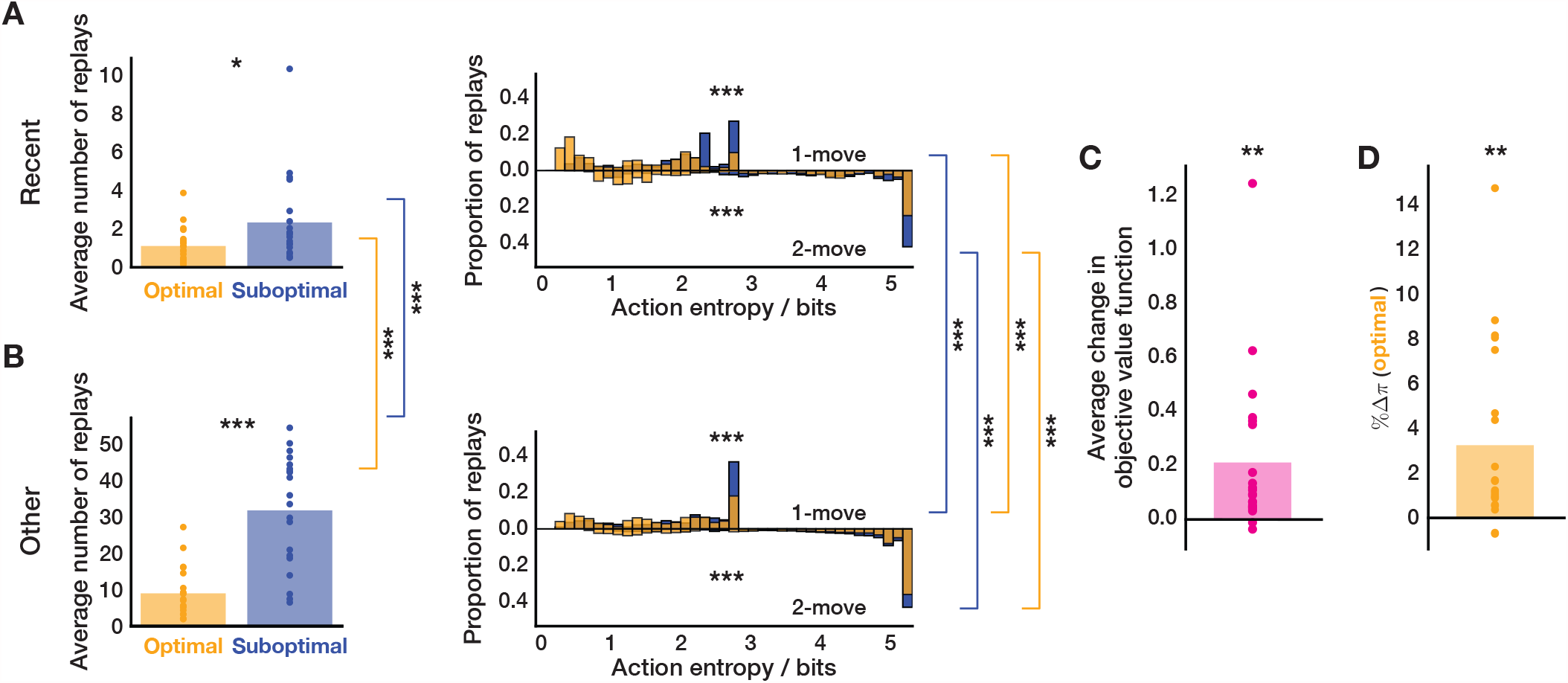
On-line Replay Statistics with Obliviated Knowledge. The layout of the figure is identical to that of Fig 6. Before engaging in off-task replay prior to blocks 3 and 5, each agent’s MF *Q*-values were zeroed-out to examine whether the overall on-line replay statistics would remain unaltered – and indeed they did. Wilcoxon rank-sum test in (A) and (B), 1-sample 2-tailed t test in (C) and (D). * *p <* 0.05, ** *p <* 0.02, *** *p <* 0.001.

**S8 Fig.**
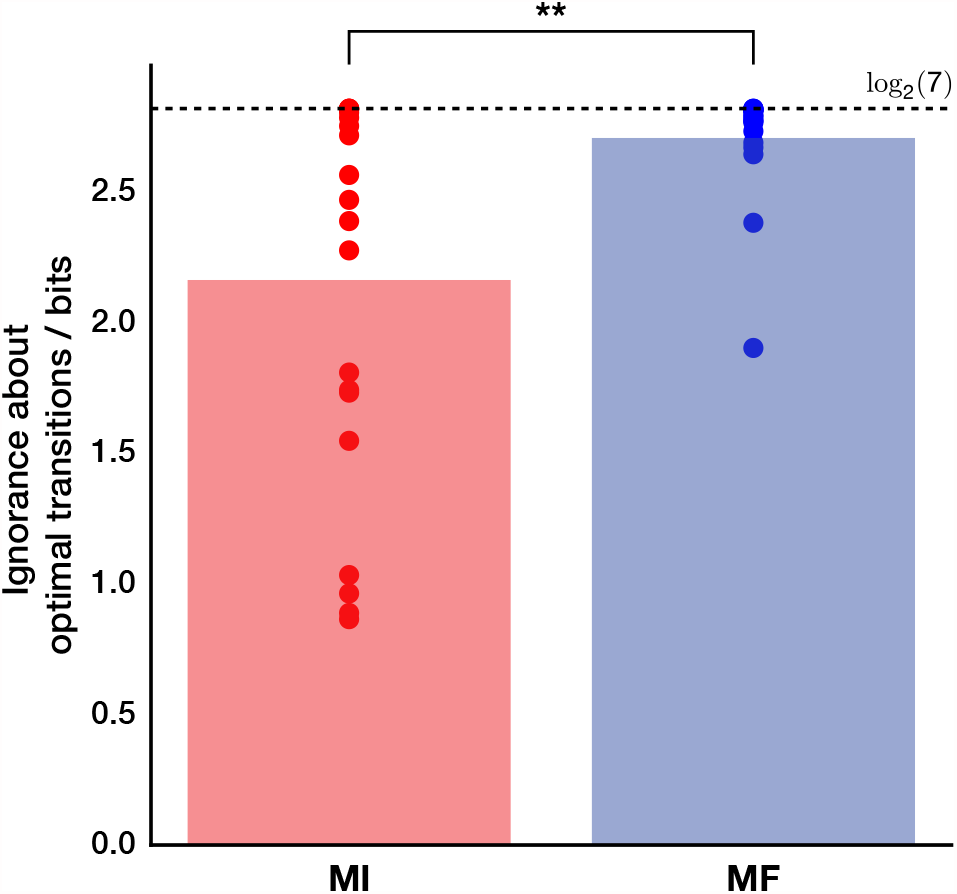
Entropy of Pre-Trained State-Transition Models. We assessed how ignorant each subject (as modelled by our agent) was about optimal 1-move transitions after learning in the training trials and immediately before entering the main task. The ignorance was computed as the average action entropy (equation 1) across all objectively optimal transitions. MI subjects who, according to our modelling, engaged in replay (*n* = 21) were significantly less ignorant about objectively optimal transitions compared to MF subjects (2-sample 2-tailed t test, *t* = −3.10, *p* = 0.003). The horizontal dashed black line shows maximum ignorance. ** *p <* 0.01.

**S9 Fig.**
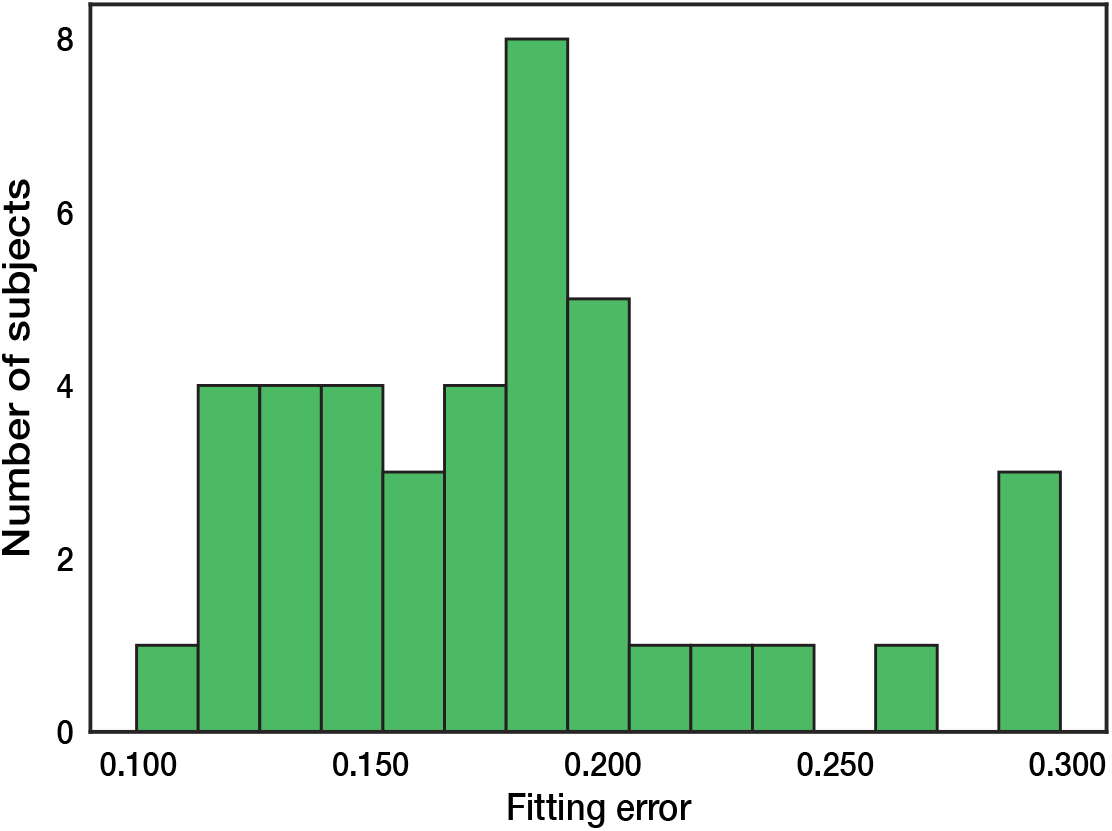
Fitting errors. Distribution of the final fitting errors (RMSD of the proportion of available reward collected between the human subjects and the agent).

## Notes

### Competing Interest Statement

The authors have declared no competing interest.

